# Targeting circPTPN12/miR-21-5p/ΔNp63α pathway as a therapeutic strategy for human endometrial fibrosis

**DOI:** 10.1101/2021.04.15.440082

**Authors:** Minmin Song, Guangfeng Zhao, Haixiang Sun, Simin Yao, Zhenhua Zhou, Peipei Jiang, Qianwen Wu, Hui Zhu, Huiyan Wang, Chenyan Dai, Jingmei Wang, Ruotian Li, Yun Cao, Haining lv, Dan Liu, Jianwu Dai, Yan Zhou, Yali Hu

## Abstract

Emerging evidence demonstrates the important role of circular RNAs (circRNAs) in regulating pathological processes in various diseases including organ fibrosis. Endometrium fibrosis is the leading cause of uterine infertility, but the role of circRNAs in its pathogenesis is largely unknown. Here, we provide the evidence that upregulation of circPTPN12 in endometrial epithelial cells (EECs) of fibrotic endometrium functions as endogenous sponge of miR-21-5p to inhibit miR-21-5p expression and activity, which in turn results in upregulation of ΔNp63α to induce the epithelial mesenchymal transition (EMT) of EECs (EEC-EMT). In a mouse model of endometrium fibrosis, circPTPN12 appears to be a cofactor of driving EEC-EMT. Our findings reveal the novel mechanism in the pathogenesis of endometrium fibrosis and the potential therapeutic strategy for endometrium fibrosis via targeting circPTPN12/miR-21-5p/ΔNp63α pathway.

## Introduction

Endometrial fibrosis is clinically characterized with intrauterine adhesions (IUA), and is often secondary to severe injury of endometrium including various uterine operations or severe inflammation. Endometrial fibrosis is the most common reason of uterine infertility (***Yu et al., 2008; March, 2011a; March, 2011b***). The mechanism of endometrial fibrosis remains unclear, which hampers the development of effective therapeutics for the disease (***March, 2011b***). Recently, our study showed that the ectopic expression of ΔNp63α, a transcription factor, in the endometria of IUA patients triggers the epithelial mesenchymal transition (EMT) of endometrial epithelial cells (EECs) (EEC-EMT) to promote endometrial fibrosis (***Zhao et al., 2020***).

In the process of EMT, post-transcriptional regulation has critical roles (***Nieto et al., 2016***). Studies showed that let-7d is downregulated in EMT of the lung epithelial cells and downregulation of let-7d in mice causes lung fibrosis (***Pandit et al., 2010***).

miR-29b suppresses EMT of lung epithelial cells and prevents mice against lung fibrosis (***Sun et al., 2019***). In the renal fibrosis, miR-34a induces EMT in renal tubular epithelial cells and knockout miR-34a ameliorates renal fibrosis in mice (***Liu et al., 2019***). Moreover, the miRNA may function differently in the pathogenesis of fibrosis in different tissues. For example, miR-27a prevents the fibrosis of detrusor and kidney (***Wu et al., 2018; Zhang et al., 2018***), but promotes the myocardial fibrosis (***Zhuang et al., 2018***). However, the role of miRNAs in the pathogenesis of endometrial fibrosis is less studied (***Li et al., 2016***).

Recent studies demonstrate that the expression and activity of miRNAs are regulated by circular RNAs (circRNAs). circRNAs are recognized as competing endogenous RNAs (ceRNAs) to sponge miRNAs to modulate the expression and activity of miRNAs and their target genes (***Li et al., 2018; Zheng et al., 2016; Cheng et al., 2019***). To date, little is known about the biological function of circRNAs in the pathogenesis of endometrial fibrosis. Since we found that EMT triggered endometrial fibrosis by the ectopic expression of ΔNp63α in EECs (***Zhao et al., 2020***), and ΔNp63α is regulated by multiple miRNAs in epidermal eratinocytes (***Candi et al., 2015; Lena et al., 2008; Rodriguez et al., 2016***), we speculated whether circRNAs is involved in EEC-EMT and in the pathogenesis of endometrial fibrosis by interacting with miRNAs. Therefore, we studied the expression changes of circRNAs/miRNAs in endometrial fibrosis and their regulatory relationship with ΔNp63α to favor the understanding of endometrial fibrosis and open promising therapeutic opportunities for IUA patients.

## Results

### miR-21-5p is downregulated in EECs and associated with EEC-EMT

To determine the transcriptomic alterations in the endometria of the patients with IUA, we performed high-throughput sequencing analysis using endometrium samples in late proliferative phase from three severe IUA patients and three normal controls. The histopathological results of the samples used for high-throughput sequencing showed endometrial fibrosis in IUA patients (***Figure 1-figure supplement 1A***). With the significance threshold of absolute mean fold change > 2 and *P* values < 0.05, of 1681 detectable miRNAs in total, 56 were upregulated and 150 were downregulated in IUA patients, compared with those in normal controls (***Figure 1A and Supplementary File 1***). These differentially expressed miRNAs were also presented in a heat map (***Figure 1-figure supplement 1B***). The most significantly increased and decreased miRNAs were verified by qRT-PCR (***Figure 1-figure supplement 1C***).

**Figure 1.**
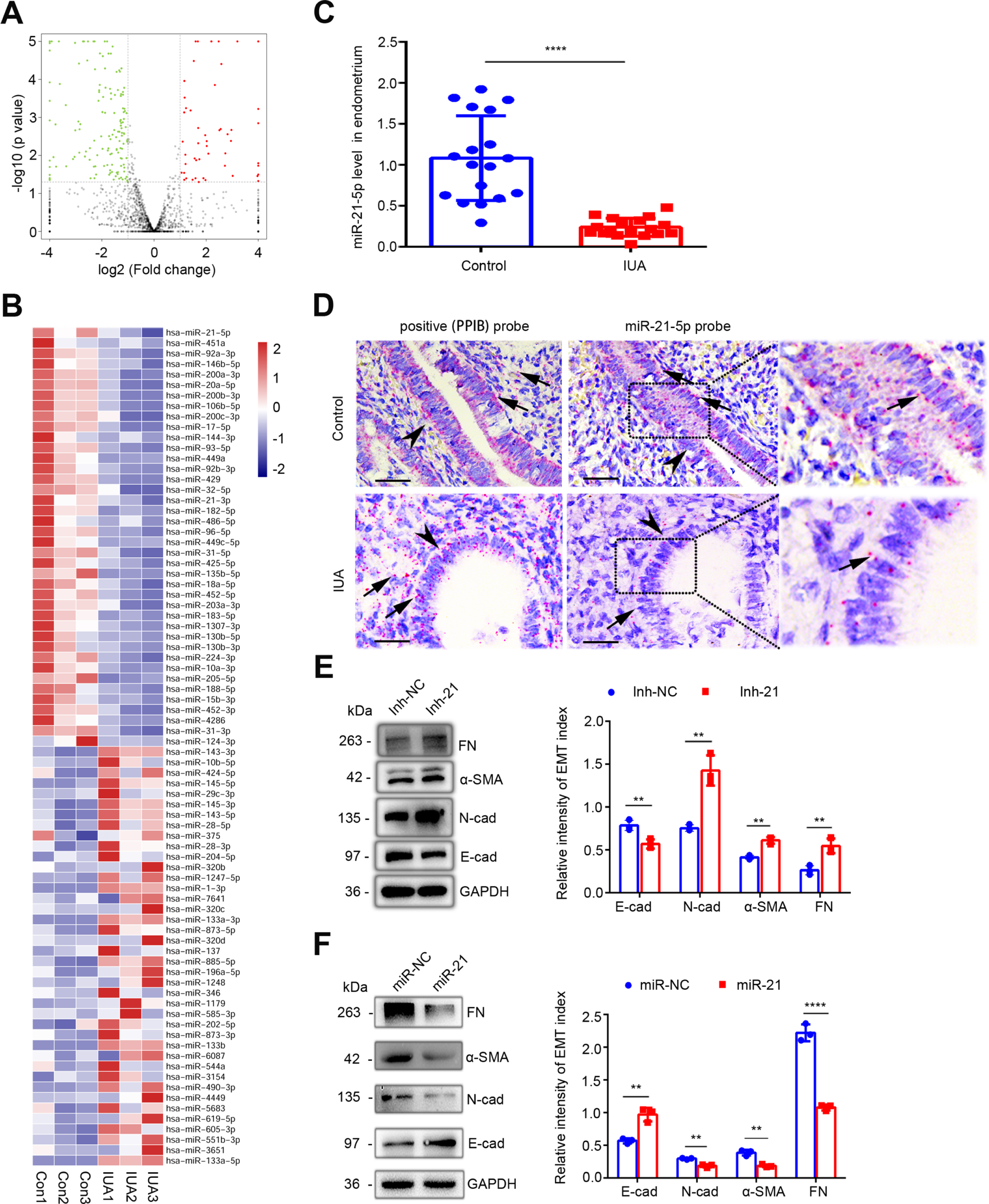
Downregulation of miR-21-5p in endometria of IUA patients promotes EEC-EMT. (**A**) Volcano plots of miRNA-sequencing of the endometria from severe IUA patients (*n* = 3) and controls (*n* = 3) based on the high-throughput RNA-sequencing analysis. The red dots represent the miRNAs up-regulated and the green ones represent the down-regulated (*P* <0.05 and fold change > 2). (**B**) A heatmap showing 40 upregulated and 40 downregulated miRNAs with high abundance in endometrium samples from severe IUA patients (*n* = 3) and controls (*n* = 3). (**C**) miR-21-5p mRNA levels in endometria of severe IUA patients (*n* = 18) and controls (*n* = 18) determined by qRT-PCR. (**D**) Representative images of RNA-scope assay using specific probes to detect miR-21-5p in the endometria of severe IUA patients (*n* = 3) and controls (*n* = 3). Peptidylprolyl isomerase B (PPIB) serves as the positive control. Arrowhead: epithelial cells; arrow (red dots): PPIB or miR-21-5p positive dots. Scale bars 50 μm. (**E**) Left: fibronectin (FN), α-smooth muscle actin (α-SMA), E-cadherin (E-cad), N-cadherin (N-cad) and GAPDH protein levels determined by Western blotting in miR-21-5p inhibitor (inh-21)- or negative control (inh-NC)-transfected (48 hours) EECs (*n* = 3). Right: The quantitative band intensities determined by image J software. (**F**) Left: FN, α-SMA, E-cad, N-cad and GAPDH protein levels determined by immunoblotting in miR-21-5p mimic (miR-21)- or negative control (miR-NC)-transfected (48 hours) EECs (*n* = 3). Right: The quantitative band intensities determined by image J software. The error bars in (C), (E) and (F) indicate mean ± SD. ** *P* < 0.01, **** *P* < 0.0001. Source data 1. Uncropped Western blots for Figure 1E. Source data 2. Uncropped Western blots for Figure 1F.

Among these dysregulated miRNAs, 40 upregulated and 40 downregulated miRNAs with higher expression abundance were shown in ***Figure 1B***. The expressive abundance of miR-21-5p was the highest in endometria of normal controls but was significantly downregulated in IUA patients (***Figure 1C and Supplementary File 2***), which was further proved by RNA-scope assay (***Figure 1D***). Meanwhile, RNA-scope assay showed that miR-21-5p was mainly expressed in EECs (***Figure 1D***). Since epithelial cells often switch to matrix-producing myofibroblasts via EMT in tissue fibrosis process (***Iwano et al., 2002; Zeisberg et al., 2007***) and we proved recently that EEC-EMT participates in endometrium fibrosis in IUA patients (***Zhao et al., 2020***), we speculated that miR-21-5p may serve as a protector for maintaining endometrium homeostasis and preventing EECs from EMT. Thus, we performed loss and gain of miR-21-5p function by transfecting miR-21-5p mimic and inhibitor into primary EECs respectively. miR-21-5p expression increased 11 times in EECs transfected with miR-21-5p mimic, while the expression decreased 2.5 times after miR-21-5p inhibitor transfection (***Figure 1-figure supplement 2***). After miR-21-5p knockdown, the expression of epithelium-specific marker, E-cadherin, was downregulated but mesenchymal markers, N-cadherin, α-smooth muscle actin (α-SMA), and fibronectin (FN), were overexpressed in EECs (***Figure 1E***). In contrast, overexpression of miR-21-5p markedly decreased the expression of N-cadherin, α-SMA, FN and increased the expression of E-cadherin (***Figure 1F***).

### circPTPN12 is upregulated in EECs and negatively correlated with miR-21-5p

Since circRNAs are the main regulators for miRNAs activity and act as ceRNA to sponge miRNAs by complementary base pairing to regulate the expression of target mRNAs (***Li et al., 2018; Zheng et al., 2016; Cheng et al., 2019***), we analyzed the expression profile of circRNAs in endometrium samples from severe IUA patients and normal controls through high-throughput sequencing. We detected 17134 distinct circRNAs and 4886 of them have been registered in circBase database (***Figure 2-figure supplement 1A***). We annotated these circRNAs and found that 4832 of them derived from protein-coding exons (***Figure 2-figure supplement 1B***). With the criteria of absolute mean fold change > 2.0 and *P* values < 0.05. The results showed that there were 263 upregulated circRNAs and 195 downregulated circRNAs (***Figure 2-figure supplement 1C***). Then, we predicted circRNAs with binding sites for miR-21-5p based on miRanda database, and found that there were 172 candidate circRNAs (***Figure 2-figure supplement 1D and Supplementary File 3***). Of the 172 miR-21-5p binding circRNAs, 12 were up-regulated and 4 were down-regulated in IUA patients (***Figure 2A and 2B***). We focused on the upregulated circRNA candidates, because miR-21-5p was downregulated in the endometria of IUA patients (***Figure 1C and 1D***) and circRNAs function as ceRNAs to inhibit miRNAs activities and expression. The most upregulated circRNA was hsa_circ_0003764 (circBase ID), and it was 64-fold higher than that in controls (***Figure 2B***). qRT-PCR (***Figure 2C***) and RNA-scope assay (***Figure 2D***) showed that hsa_circ_0003764 was significantly upregulated in the endometria of IUA patients and mainly expressed in EECs.

**Figure 2.**
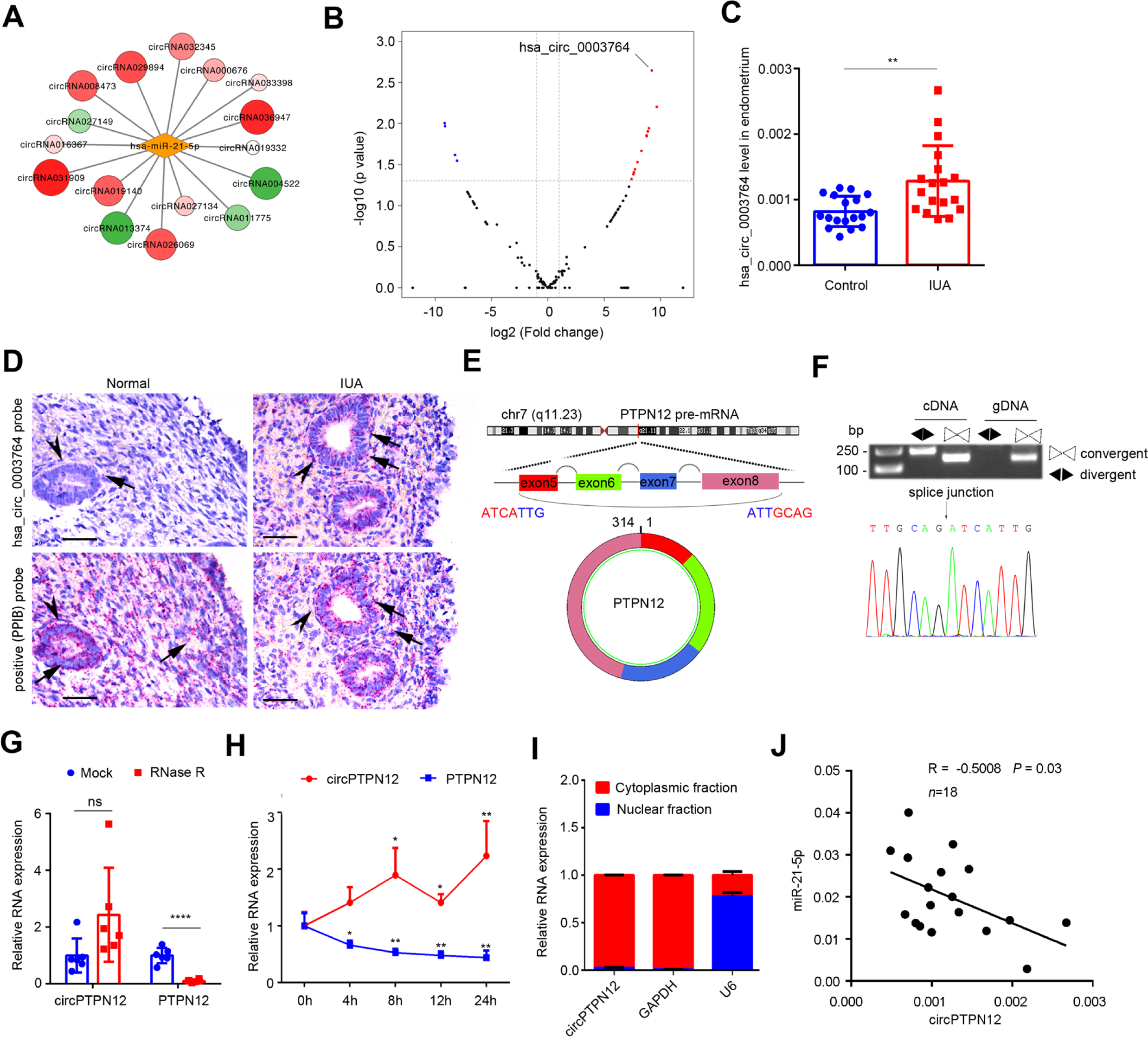
Upregulation of circPTPN12 in endometria of IUA patients is negatively correlated with miR-21-5p. (**A**) Network diagram of circRNAs that have binding sites for miR-21-5p and that significantly express in the endometria from severe IUA patients (*n* = 3) and controls (*n* = 3) based on the high-throughput RNA-sequencing analysis. (**B**) Volcano plots of circRNAs possessing binding sites for miR-21-5p in the endometria from severe IUA patients (*n* = 3) and controls (*n* = 3) based on the high-throughput RNA-sequencing analysis. The red dots represent the circRNAs up-regulated and the blue dots represent the circRNAs down-regulated (*P* < 0.05 and fold change > 2). (**C**) Hsa_circ_0003764 expression levels in endometria of severe IUA patients (*n* = 18) and controls (*n* = 18) determined by qRT-PCR. (**D**) Representative images of RNA-scope assay using specific probes to detect hsa_circ_0003764 in the endometria of severe IUA patients (*n* = 3) and controls (*n* = 3). PPIB serves as the positive control. Arrowhead: epithelial cells; arrow (red dots): PPIB or has_circ_0003764 positive dots. Scale bars 50 μm. (**E**) Schematic illustration of the genomic loci of tyrosine-protein phosphatase non-receptor type 12 (PTPN12) gene and the cyclization of circPTPN12. (**F**) Verification of circPTPN12 in endometria. Top: Agarose gel electrophoresis shows that divergent primers amplified circPTPN12 in complementary DNA (cDNA) but not in genomic DNA (gDNA). Bottom: Sanger sequencing of the amplified band with divergent primers shows the spliced junction of circPTPN12. (**G**) qRT-PCR analysis of circPTPN12 and PTPN12 mRNA levels in total RNA extracted from endometria of severe IUA patients (*n* = 6) with or without RNase R treatment. (**H**) qRT-PCR analysis of circPTPN12 and PTPN12 mRNA levels in EECs (*n* = 4) treated with actinomycin D at the indicated time points. (**I**) circPTPN12 expression level in the nuclear and cytoplasmic fractions of EECs (*n* = 4) determined by qRT-PCR. (**J**) The correlation of circPTPN12 and miR-21-5p in endometria of IUA patients (*n* = 18). Spearman’s correlation coefficient R = –0.5008, *P* = 0.03. (C), (G) and (H) Error bars indicate mean ± SD. No statistical difference (ns), * *P* < 0.05, ** *P* < 0.01, **** *P* < 0.0001. **Source data 1.** Uncropped gels for Figure 2F.

Through circPrimer software, we found that hsa_circ_0003764 is derived from 5 to 8 exons of tyrosine-protein phosphatase non-receptor type 12 (PTPN12) gene.

Therefore, we termed it circPTPN12 in following experiments (***Figure 2E***).

We designed specific divergent primers to prove the existence of circPTPN12 in the endometria of IUA patients by PCR. The predicted spliced junction of circPTPN12 was validated with agarose gel electrophoresis (***Figure 2F Top***). The resultant products of divergent primers were confirmed in line with the sequence of circPTPN12 by sequencing (***Figure 2F Bottom***). To investigate the stability of circPTPN12, total RNA isolated from IUA patients’ endometria was treated with RNase R for 15 minutes, then the expression of circPTPN12 and PTPN12 was respectively detected and the results showed that PTPN12 expression was significantly decreased while circPTPN12 expression was not changed (***Figure 2G***). Furthermore, we added actinomycin D to inhibit the transcription of EECs. We harvested total RNA at five time points, analyzed the expression of circPTPN12 and PTPN12 by qRT-PCR, and revealed that the half-life of circPTPN12 was exceeding 24 hours, whereas the half-life of linear PTPN12 transcript was less than 4 hours (***Figure 2H***). Thus, circPTPN12 was more stable than PTPN12 mRNA, suggesting that circPTPN12 is circular. Separation analysis of nuclear and cytoplasm showed that circPTPN12 was mainly distributed in the cytoplasm of EECs (***Figure 2I***).

Since miR-21-5p was decreased and circPTPN12 was increased in fibrotic endometria of IUA patients, and both were expressed in EECs, we further analyzed the correlation between them. The results showed that circPTPN12 was negatively correlated with miR-21-5p (***Figure 2J***).

### circPTPN12 functions as ceRNA to sponge miR-21-5p

To prove the interaction between circPTPN12 and miR-21-5p, we constructed a circPTPN12 plasmid to transfect HEK-293T cells. The transfection efficiency was verified by qRT-PCR (***Figure 3A***). We used biotin-labeled circPTPN12 to pull down the circPTPN12 in the lysates of the transfected HEK-293T cells. The electrophoresis showed that miR-21-5p was pulled down by circPTPN12 (***Figure 3B***). To determine the binding site of circPTPN12 for miR-21-5p, we cloned the full-length sequence of circPTPN12 in pGL3 vector containing a luciferase reporter. Luciferase activity was substantially reduced in the presence of the miR-21-5p mimic, which was effectively restored when miR-21-5p was knockdown (***Figure 3C***). We then compared the sequence of circPTPN12 with that of miR-21-5p using circBank and Circular RNA Interactome databases and noticed that circPTPN12 contains three putative target sites of miR-21-5p (***Figure 3D Top***). The test of delete mutations of each or all of three putative binding sites showed that only the third site was validated to have the capacity of binding by the luciferase assay (***Figure 3D Bottom***). The schematic diagram of real binding site is shown in ***Figure 3E***. To further verify the binding of circPTPN12 with miR-21-5p, we conducted RNA immunoprecipitation (RIP) experiment to precipitate the complex of argonaute (AGO) protein based on a principle that miRNAs exert post-transcriptional regulation through binding with AGO protein to form the targeting module of the miRNA-induced silencing complex (miRISC) (***Gebert et al., 2019***). The RIP experiment showed that the precipitates of HEK-293T cell lysates stably expressing Flag-AGO2 or Flag-GFP contained not only miR-21-5p, but also circPTPN12 (***Figure 3F***), indicating the bind of circPTPN12 and miR-21-5p. Fluorescence in situ hybridization (FISH) demonstrated that miR-21-5p was colocalized with circPTPN12 in the cytoplasm of Ishikawa cell (***Figure 3G***).

**Figure 3.**
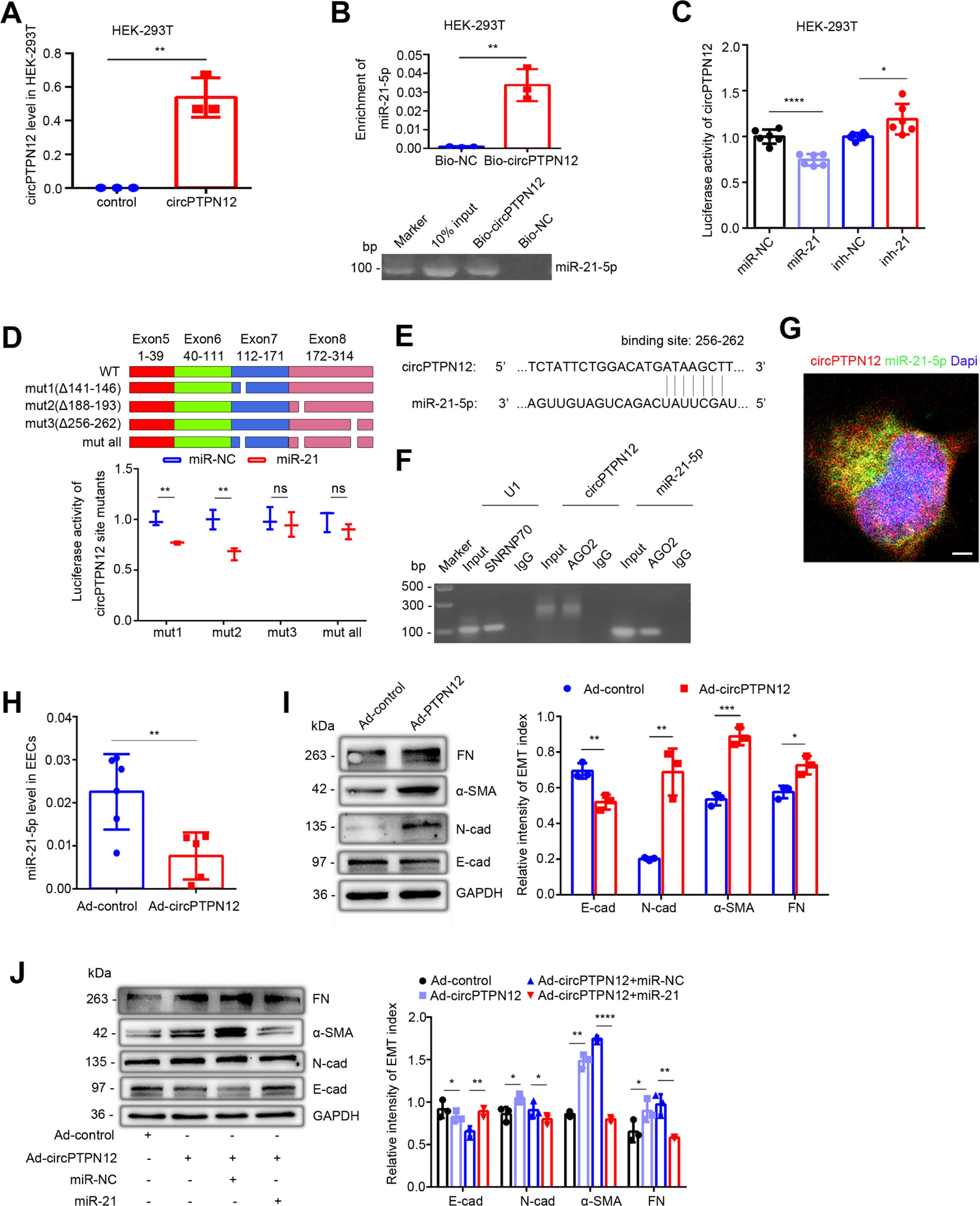
circPTPN12 functions as ceRNA to sponge miR-21-5p. (**A**) qRT-PCR analysis of circPTPN12 expression level in circPTPN12 plasmid transfected (24 hours) HEK-293T cells (*n* = 3). (**B**) Capture of miR-21-5p by circPTPN12. Top: qRT-PCR analysis of miR-21-5p pulled down by biotin-labeled circPTPN12 (Bio-circPTPN12) or scramble (Bio-NC) probe from the HEK-293T cell lysates transfection with circPTPN12 plasmid (*n* = 3). Bottom: Amplified qRT-PCR products in agarose gel electrophoresis. (**C**) Luciferase activity of circPTPN12 in HEK-293T cells transfected with miR-21-5p mimic (*n* = 6) or inhibitor (*n* = 6). (**D**) Top: Schematic diagram of three mutated sites in circPTPN12 luciferase reporter gene. Bottom: Luciferase activity of Luc-circPTPN12 containing single or all mutated sites of three putative miR-21-5p binding sites in HEK-293T cells transfected with miR-21-5p mimic (*n* = 3). (**E**) Simulated diagram of the exact binding sites between circPTPN12 and miR-21-5p. (**F**) Enrichment of circPTPN12 and miR-21-5p in AGO2 immunoprecipitates of HEK-293T cells transfected with circPTPN12 plasmid in agarose gel electrophoresis. (**G**) Co-localization between circPTPN12 and miR-21-5p by RNA FISH in Ishikawa cells. Nuclei were stained with DAPI. Scale bar, 1 μm. (**H**) qRT–PCR analysis of miR-21-5p level in adenovirus containing circPTPN12 (Ad-circPTPN12) or adenovirus containing no circPTPN12 (Ad-control) infected (48 hours) EECs (*n* = 5). (**I**) Left: FN, α-SMA, N-cad, E-cad, and GAPDH protein levels determined by Western blotting in circPTPN12-infected (72 hours) EECs (*n* = 3). Right: The quantitative band intensities determined by image J software. (**J**) Left: FN, α-SMA, N-cad, E-cad, and GAPDH protein levels determined by Western blotting in EECs transfected with miR-21-5p mimic or miR-NC for 48 hours in the presence of Ad-circPTPN12 (*n* = 3). Right: The quantitative band intensities determined by image J software. (A)-(D) and (H)-(J) Error bars indicate mean ± SD. No statistical difference (ns), * *P* < 0.05, ** *P* < 0.01, *** *P* < 0.001, **** *P* < 0.0001. **Source data 1.** Uncropped Western blots for Figure 3I. **Source data 2.** Uncropped Western blots for Figure 3J.

To investigate the effect of circPTPN12 on the function of miR-21-5p, we constructed recombinant adenovirus harboring exons 5-8 of PTPN12 (Ad-circPTPN12) along with approximately 1 kb flanking intron sequences containing complementary Alu elements (***Figure 3-figure supplement 1A***). The expression of circPTPN12 increased 7 times in EECs after Ad-circPTPN12 infection (***Figure 3-figure supplement 1B***).

Importantly, qRT-PCR confirmed that circPTPN12 overexpression decreased the miR-21-5p level in primary EECs (***Figure 3H***). Meanwhile, compared with that of adenovirus containing no circPTPN12 (Ad-control), overexpression of circPTPN12 enhanced protein abundance of N-cadherin, α-SMA, and FN, but decreased the level of E-cadherin in EECs (***Figure 3I***). While re-transfected with miR-21-5p mimic reversed this phenomenon with upregulated E-cadherin expression, and downregulated N-cadherin, α-SMA and FN expression (***Figure 3J***).

### Downregulation of miR-21-5p by circPTPN12 promotes EEC-EMT through upregulating ΔNp63α

Since circPTPN12 inhibited miR-21-5p activity to promote EEC-EMT, we tried to identify its target gene(s). Our recent study showed that ΔNp63α induces EEC-EMT in IUA patients (***Zhao et al., 2020***). In the present study, we found that 3 ’UTR of ΔNp63α has miR-21-5p binding sites by prediction in Microrna.org database.

Therefore, we conducted the luciferase reporter assay with the full length 3’-UTR of ΔNp63α^wild^ and ΔNp63α^mut^ targeted by miR-21-5p (***Figure 4A***). In line with the prediction, transfection with miR-21-5p mimic significantly reduced luciferase activity in HEK-293T cells transfected with ΔNp63α^wild^ plasmid, whereas knockout of miR-21-5p in ΔNp63α^wild^ HEK-293T cells enhanced the luciferase activity; however, these effects were not observed in HEK-293T cells transfected with ΔNp63α^mut^ plasmid (***Figure 4B***). We used primary EECs transfected with the miR-21-mimic or inhibitor for 48 hours following infection with recombinant adenovirus containing ΔNp63α (Ad-ΔNp63α) and empty vector (Ad-CTL) respectively, and found that miR-21-5p mimic remarkably downregulated the mRNA and protein of ΔNp63α in ΔNp63α+ EECs (***Figure 4C and 4D***). And the miR-21-5p inhibitor upregulated ΔNp63α protein level in ΔNp63α+ EECs (***Figure 4E***). Meanwhile, upregulation of circPTPN12 increased luciferase activity in HEK-293T cells transfected with ΔNp63α^wild^ plasmids, while no luciferase activity change was observed when HEK-293T transfected with ΔNp63α^mut^ plasmids (***Figure 4F***). Moreover, the upregulation of circPTPN12 counteracted the inhibitory effect of miR-21-5p on ΔNp63α expression (***Figure 4G***). We also revealed that miR-21-5p mimic remitted ΔNp63α-induced EEC-EMT, upregulated E-cadherin and downregulated N-cadherin and α-SMA at both the mRNA and protein levels (***Figure 4H and 4I***). Furthermore, upregulation of circPTPN12 counteracted the reversal effect of miR-21-5p on ΔNp63α-induced EMT in EECs (***Figure 4J***).

**Figure 4.**
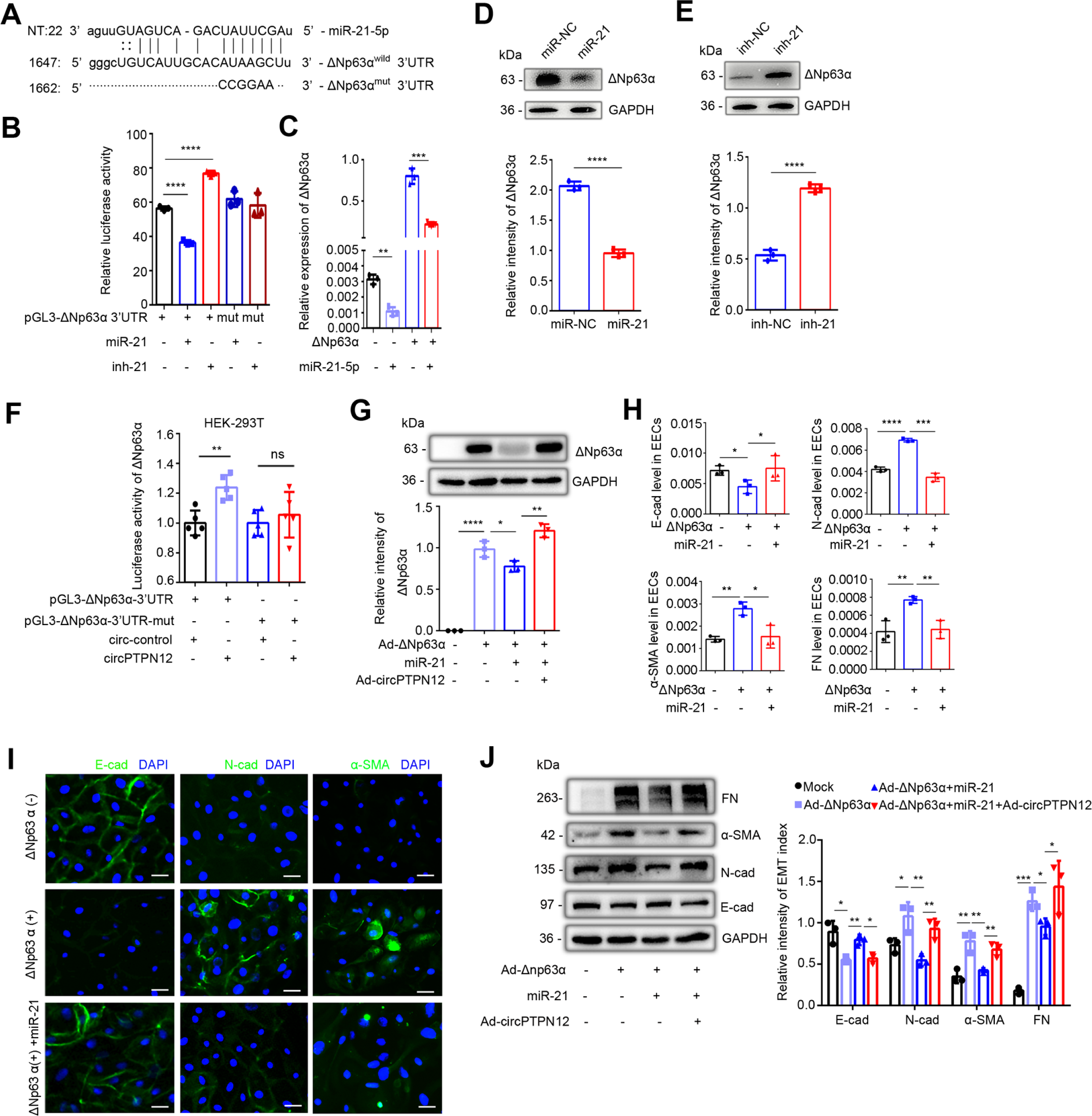
circPTPN12-downregulated miR-21-5p promotes EEC-EMT through upregulation of ΔNp63α. (**A**) The putative site for the interaction between miR-21-5p and the 3’UTR of ΔNp63α. The predicted miR-21-5p binding sequence AUAAGC was replaced by CCGGAA for pGL3-ΔNp63α^mut^ construction. (**B**) Luciferase activity of ΔNp63α^wild^ and ΔNp63α^mut^ in HEK-293T cells (*n* = 3) co-transfected with miR-21-5p mimic or inhibitor for 24 hours. (**C**) qRT-PCR analysis of ΔNp63α levels in adenovirus containing ΔNp63α (Ad-ΔNp63α) infected EECs transfected with miR-21-5p mimic (*n* = 3). (**D**) Top: ΔNp63α protein expression in miR-21-5p mimic transfected-ΔNp63α+ EECs (48 hours) determined by Western blotting (*n* = 3). Bottom: The quantitative band intensities determined by image J software. (**E**) Top: ΔNp63α protein expression in miR-21-5p inhibitor transfected-ΔNp63α+ EECs (48 hours) determined by Western blotting (*n* = 3). Bottom: The quantitative band intensities determined by image J software. (**F**) Luciferase activity of ΔNp63α^wild^ and ΔNp63α^mut^ in HEK-293T cells co-transfected with circPTPN12 or circ-control plasmids (*n* = 5). (**G**) Top: ΔNp63α protein expression determined by Western blotting in in Ad-circPTPN12 infected ΔNp63α+ or ΔNp63α-EECs in the presence of miR-21-5p mimic or miR-NC for 48 hours (*n* = 3). Bottom: The quantitative band intensities determined by image J software. (**H**) qRT-PCR analysis of E-cad, N-cad, α-SMA and FN mRNA levels in miR-21-5p mimic or miR-NC transfected ΔNp63α+ EECs (*n* = 3). (**I**) Representative images (*n* = 3) of E-cad, N-cad and α-SMA immunofluorescence staining in miR-21-5p mimic or miR-NC transfected ΔNp63α+ EECs. Scale bars, 20 μm. (**J**) Left: FN, α-SMA, N-cad, E-cad, and GAPDH protein levels determined by Western blotting in Ad-circPTPN12 infected ΔNp63α+ or ΔNp63α-EECs in the presence of miR-21-5p mimic or miR-NC for 48 hours (*n = 3*). Right: The quantitative band intensities determined by image J software. (B)-(H) and (J) Error bars indicate mean ± SD. * *P* < 0.05, ** *P* < 0.01, *** *P* < 0.001, **** *P* < 0.0001. **Source data 1.** Uncropped Western blots for Figure 4D and Figure 4E. **Source data 2.** Uncropped Western blots for Figure 4G. **Source data 3.** Uncropped Western blots for Figure 4J.

### miR-21-5p alleviates circPTPN12 induced EEC-EMT in IUA-like mouse model

We developed a mouse model by uterine mechanical injury to simulate IUA in human as described previously (***Zhao et al., 2020; Yang et al., 2017***). Compared with sham-operated mice, the mechanically injured mice had mildly upregulated N-cadherin and α-SMA in the luminal epithelial cells of endometria, slightly downregulated E-cadherin in luminal and glandular EECs, and minimal changes in Masson staining (***Figure 5-figure supplement 1A***).

**Figure 5.**
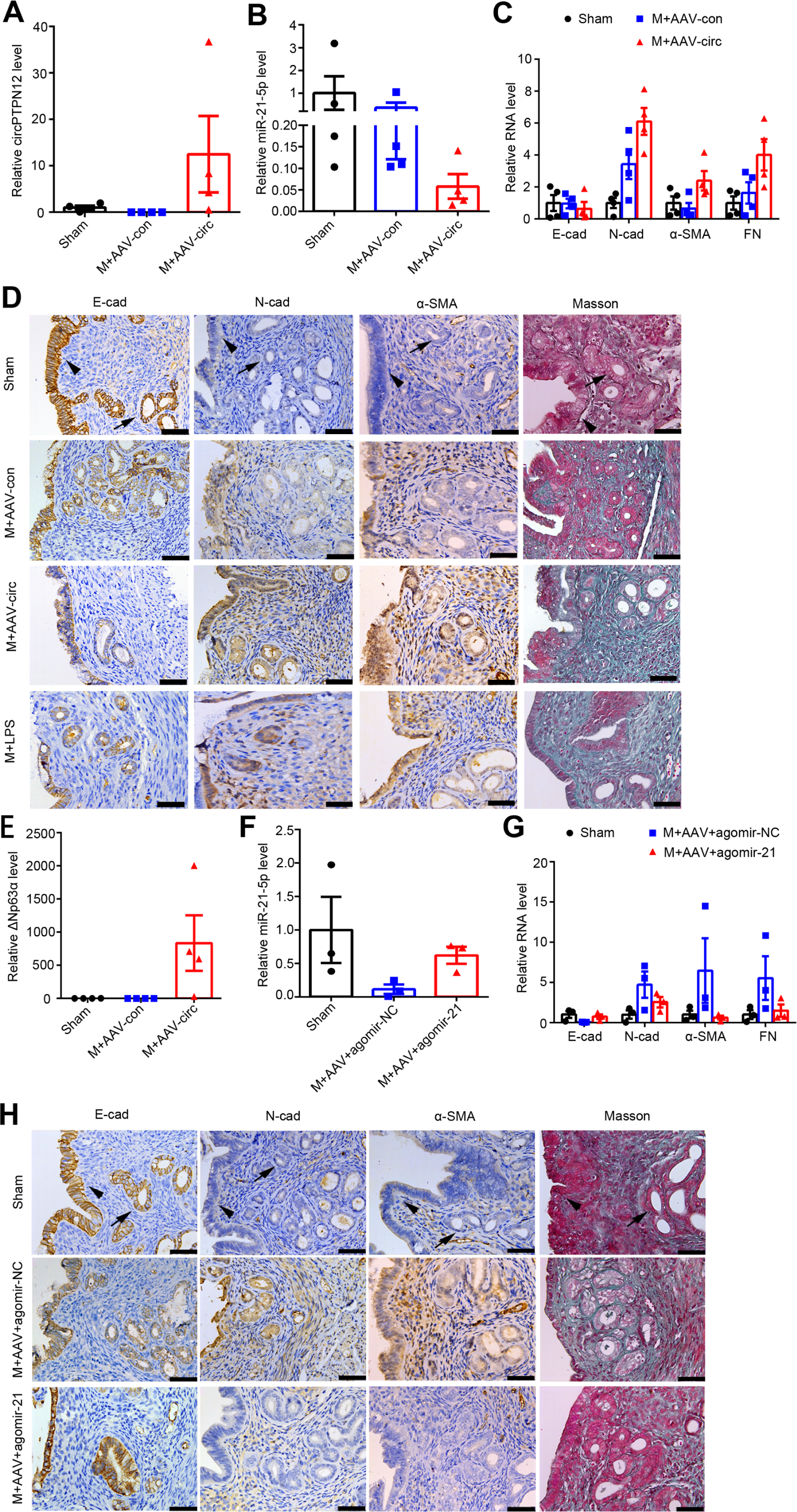
Intrauterine injection of miR-21-5p alleviates circPTPN12-induced EEC-EMT in vivo. (A)-(E) qRT-PCR analysis of circPTPN12 levels (**A**), miR-21-5p levels (**B**), E-cad, N-cad, α-SMA and FN mRNA levels (**C**), representative immunohistochemistry images of E-cad, N-cad, α-SMA and Masson staining (**D** The first three rows), ΔNp63α levels (**E**) in endometria of mice with sham-operation (Sham, *n* = 4), mechanical injury and adeno-associated virus (AAV)-control injection (M + AAV-con, *n* = 4), mechanical injury and AAV-circPTPN12 injection (M + AAV-circ, *n* = 4). (D The forth row) Representative immunohistochemistry images of E-cad, N-cad, α-SMA and Masson staining in endometria of mice with mechanical injury and lipopolysaccharide injection (M + LPS, *n* = 4). (F)-(H) qRT-PCR analysis of miR-21-5p levels (**F**), E-cad, N-cad, α-SMA and FN mRNA levels (**G**), representative immunohistochemistry images of E-cad, N-cad, α-SMA and Masson staining (**H**) in endometria of mice with sham-operation (*n* = 3), mechanical injury with AAV-circPTPN12 and agomir-NC injection (M+ AAV + agomiR-NC, *n* = 3), mechanical injury with AAV-circPTPN12 and agomir-21-5p injection (M+ AAV + agomiR-21, *n* = 3). Scale bars, 50 μm. Arrow head: luminal epithelial cells; arrow: glandular epithelial cells. (A)-(C), (E)-(F) and (G) Error bars indicate mean ± SEM.

Based on circBase and circBank database, mouse does not express circPTPN12, which provided us an opportunity to test the role of circPTPN12 in the pathogenesis of endometrium fibrosis by intrauterine injection of a circPTPN12-containing recombinant adeno-associated virus with a green label (AAV-circPTPN12). The flowchart of intrauterine injection is presented in ***Figure 5-figure supplement 1B***. The green fluorescence in mouse endometria was clearly observed after four estrous cycles following virus injection (***Figure 5-figure supplement 1C***), indicating that AAV-circPTPN12 infected the mouse endometrium. We further validated the increased circPTPN12 levels in the endometria by qRT-PCR (***Figure 5A***). With the increase of circPTPN12 expression, the expression of miR-21-5p decreased significantly (***Figure 5B***). Furthermore, the mice injected with AAV-circPTPN12 showed increased mRNA levels of N-cadherin, α-SMA and FN, and decreased mRNA level of E-cadherin, compared with those injected with adeno-associated virus with empty vector (***Figure 5C***). Immunohistochemical staining displayed that AAV-circPTPN12 injection clearly reduced E-cadherin expression in EECs and remarkably upregulated N-cadherin and α-SMA in both epithelial and stroma cells (***Figure 5D***). In addition, Masson staining showed strong positive (***Figure 5D***). The results of EMT and severe endometrial fibrosis in mice caused by AAV-circPTPN12 intrauterine injection were similar to the results induced by dual-injury, namely mechanical injury with lipopolysaccharide (LPS) injection (***Zhao et al., 2020***). ΔNp63α expression was also significantly upregulated after AAV-circPTPN12 administration (***Figure 5E***).

Since circPTPN12 served as the ceRNA of miR-21-5p and the pro-EMT effect of circPTPN12 was reversed by miR-21-5p in vitro (***Figure 3B-3I***), we further conducted the mouse experiment by injecting agomir-21-5p or agomir negative control (agomir-NC) into the uterine cavity of the AAV-circPTPN12 mouse model every five days for three times (***Figure 5-figure supplement 1D and 1E***). The results showed that miR-21-5p in the endometria was overexpressed (***Figure 5F***) and the mRNA level of E-cadherin was elevated, while the mRNA levels of N-cadherin, α-SMA and FN were decreased (***Figure 5G***). Immunohistochemical results showed that agomir-21-5p application significantly increased E-cadherin, and reducedN-cadherin and α-SMA in both luminal and glandular epithelia and reduced Masson staining compared with agomir-NC injection (***Figure 5H***).

## Discussion

In the present study, we revealed that miR-21-5p was highly expressed in EECs of normal endometrium and significantly downregulated in fibrotic endometrium in IUA patients, and that circPTPN12 was highly expressed in EECs in the patients, but was minimally expressed in normal endometrium. We further demonstrated that circPTPN12 sponged miR-21-5p, resulting in the upregulation of ΔNp63α, which promoted EEC-EMT and endometrium fibrosis. These novel findings highlight the role of interaction of circRNAs and miRNAs in the pathogenesis of endometrium fibrosis. Moreover, in the mouse model of IUA induced by mechanical injury and circPTPN12, administration of miR-21-5p in uterine cavity greatly improved endometrium fibrosis. Therefore, the present study provides potential therapeutics for IUA.

miRNAs play an important role in EMT and fibrosis processes in lung and kidney fibrosis (***Nieto et al., 2016***). Just based on the findings that miR-29b (***Sun et al., 2019***) and miR-326 (***Ning et al., 2018***) alleviated lung fibrosis, other scholars showed that miR-29b prevented endometrial fibrosis in rats (***Li et al., 2016***), and miR-326 inhibited the transition of human endometrial stromal cells to myofibroblasts in vitro (***Das et al., 2014***). However, increasing evidence has demonstrated that sufficient abundance of miRNAs transcripts in cells is essential in the regulating target genes (***Liu et al., 2019; Brown et al., 2009***). Thus, in this study we chose the candidate miRNAs not only based on their expressive difference but also based on their expressive abundance. With RNA-seq, we found that miR-21-5p was the most abundantly expressed in normal endometrium but significantly decreased in fibrotic endometria of IUA patients (***Figure 1C and 1D***). The results suggest that miR-21-5p in endometria may play a contrast role in the fibrogenesis of lung and renal (***Chau et al., 2012***), since miR-21-5p is minimally expressed in normal lung and kidney epithelial cells and highly expressed in the process of EMT in lung and kidney (***Yamada et al., 2013; Wang et al., 2019***). Loss-and-gain test showed that loss of miR-21-5p resulted in remarkable decrement of epithelial markers and substantial increment of mesenchymal markers (***Figure 1E***), and gain of miR-21-5p reversed these phenomena (***Figure 1F***). These results indicate that miR-21-5p is essential for maintaining EEC homeostasis and downregulation of miR-21-5p could induce EEC-EMT.

Having demonstrated the essential role of miR-21-5p in the homeostasis of EEC and the downregulation of miR-21-5p in IUA patients, we tried to figure out the regulatory factor(s) of miR-21-5p. We analyzed circRNA profile (***Figure 2A and 2B***), and the results showed that circPTPN12 was significantly upregulated in endometria of IUA patients, whereas it was rarely expressed in endometria of normal controls (***Figure 2C***). RNA scope demonstrated that circPTPN12 was located in EEC (***Figure 2D***), and further Nucleo-cytoplasmic separation experiment showed that circPTPN12 was localized in the cytoplasm of EECs (***Figure 2I***). FISH results showed that both circPTPN12 and miR-21-5p were co-localized in the cytoplasm (***Figure 3G***).

Bioinformatic analysis indicated that circPTPN12 has binding sites for miR-21-5p (***Figure 2A and Figure 3D***), which was validated by luciferase activities (***Figure 3C and 3D***). Moreover, circPTPN12 interacted with miR-21-5p in RNA pull-down and RIP experiments (***Figure 3B and 3F***), leading to reduced expression of miR-21-5p in EECs (***Figure 3H***), which further decreased the level of the epithelial marker and increased level of mesenchymal markers (***Figure 3H***). These results indicate that circPTPN12 functions as miR-21-5p sponge.

Previously, we reported that ΔNp63α is increased in EECs of fibrotic endometria in IUA patients and the inhibition of ΔNp63α level in EECs alleviates EEC-EMT in cell culture and in experimental model (***Zhao et al., 2020***). In the present study, we revealed that downregulation of miR-21-5p increased ΔNp63α level in EECs (***Figure 4E***), followed by EEC-EMT (***Figure 4H and 4I***). The results indicate that ΔNp63α is regulated by miR-21-5p.

The establishment of mouse IUA models usually require dual-injury, mechanical injury and LPS (***Zhao et al., 2020; Yao et al., 2019***). In the present study, the mice with mechanical injury alone only showed mild EEC-EMT tendency (***Figure 5-figure supplement 1A***), whereas the mice with mechanical injury and intrauterine injection of circPTPN12 displayed typical EEC-EMT and fibrosis (***Figure 5C and 5D***), which was similar to those in the mice with mechanical injury and LPS treatment (***Zhao et al., 2020***). Furthermore, replenishing miR-21-5p by intrauterine injection in mouse modal offset the pro-EMT effect of circPTPN12 on EECs (***Figure 5G and 5H***). These results indicated the role of circPTPN12 in the pathogenesis of IUA and the potential therapeutic effect of miR-21-5p in IUA patients.

There are some limitations in our study. The endometrial injuries in all IUA patients were caused by repeated curettage. Thus, it is unknown whether the results found in the present study are applicable to IUA patients caused by other factors such as tuberculosis or uterine artery embolization. Another limitation is that we used mouse IUA-like model only. For the potential clinical application in future, more studies are required to confirm the therapeutic effects in other animal models such as rats, pigs, and non-human primates.

Based on the aforementioned results, we propose following working model (***Figure 6***).

**Figure 6.**
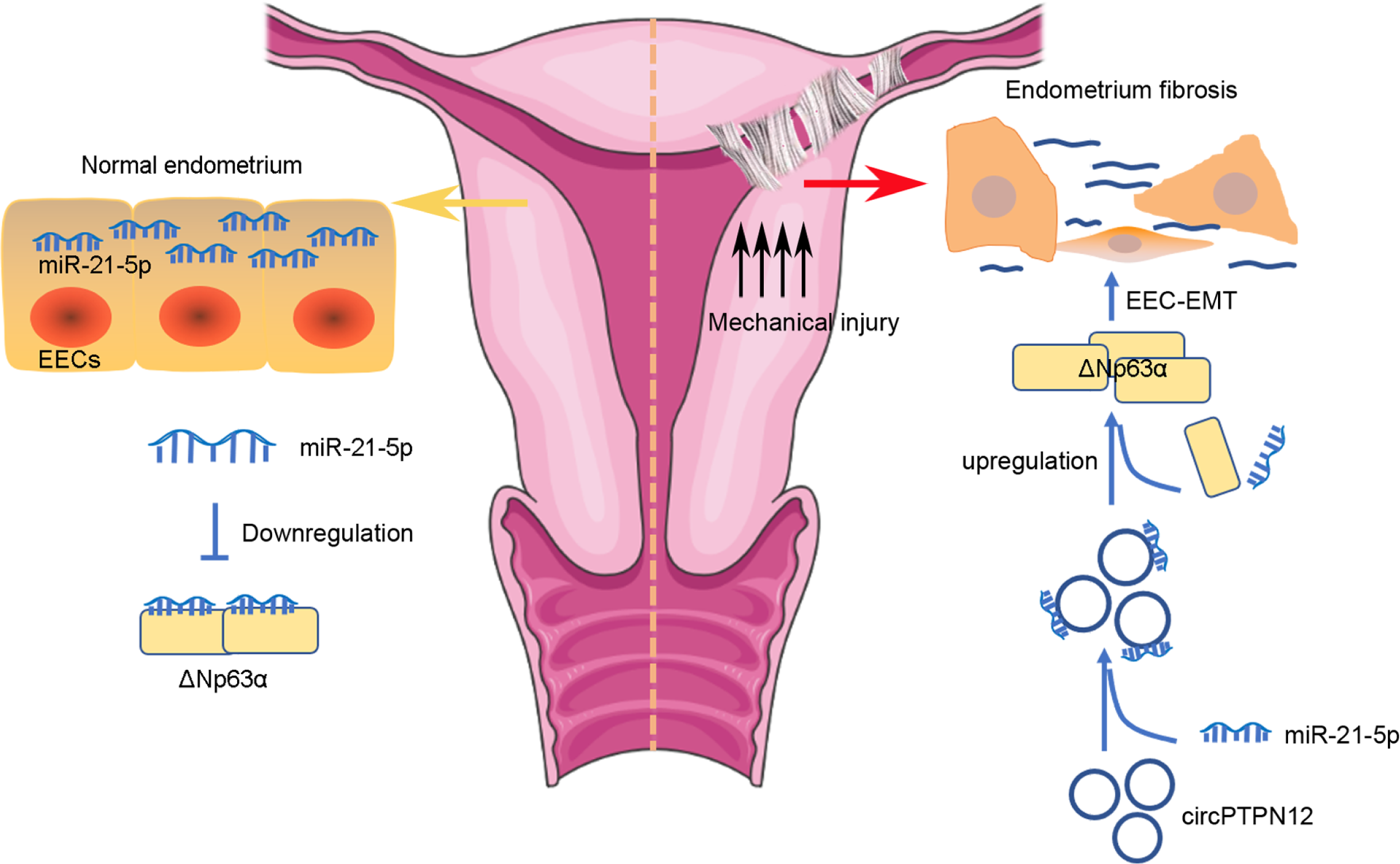
Proposed pathogenesis of endometrium fibrosis.

In normal endometrium, the high level of miR-21-5p inhibits ΔNp63α level in EECs to maintain the homeostasis of EECs. In the pathological circumstance, circPTPN12 is highly expressed, which decreases miR-21-5p level in EECs, so that the suppression effect of miR-21-5p on the expression of ΔNp63α is counteracted, leading to EEC-EMT and endometrium fibrosis. Thus, we uncovered a novel mechanism in the development of IUA. Our findings suggest that circPTPN12/miR-21-5p/ΔNp63α pathway may be a novel therapeutic target for IUA.

## Materials and methods

### Human endometrium samples

This study was approved by the Committee on Human Research of the Nanjing Drum Tower Hospital (No. 2012022), and informed consent was obtained from each participant. Human endometrium samples were collected in the late proliferative phase of the menstrual cycle from child-bearing age women during hysteroscopy for infertility screening at the Affiliated Drum Tower Hospital of Nanjing University from January 2014 to December 2016. The late proliferative phase was defined based on follicle size between 15 to 18 mm by ultrasonography and a low level of serum progesterone. This study enrolled 21 normal controls, who were infertile patients with normal ovary function, normal uterine cavity and endometrium thickness ≥ 8 mm immediate before ovulation. Twenty-one severe IUA patients were diagnosed as score >8 based on criteria recommended by the American Fertility Society (***American Fertility Society, 1988***). The clinical information of all patients and controls was listed in ***Supplementary File 4***. Fibrotic endometrial samples were taken from two sites in the uterine body or fundus that showed the most severe adhesion. Normal control samples were taken in the uterine body and fundus. Half of each sample was stored in liquid nitrogen and the other half was fixed with formalin and used for further experiments as described below.

### RNA sequencing, bioinformatics analysis for miRNAs and circRNAs

Total RNA extracted from the endometrial samples was subjected to library construction, followed by sequenced on an Illumina Hiseq 250020 (miRNA) or Illumina Hiseq X10 platform (circRNA) by Vazyme (Nanjing, China). The raw reads were filtered to achieve the clean tags through in-house Perl processes to map to the reference genome. miRNAs were identified by mapping to miRBase database and circRNAs were identified by circRNA Finder database (***Fu et al., 2014***). The miRNAs and circRNAs expression level were calculated and normalized to transcripts per million (TPM), TPM = 10^6 C/L, where C is the counts of a miRNA or circRNA, and L is counts sum of all miRNAs or circRNAs. To identify differentially expressed miRNAs and circRNAs between samples, the edgeR package (http://www.r-project.org/) was used. Differentially expressed miRNAs and circRNAs were identified with fold change > 2 or < −2 and *p* < 0.05.

### Cell isolation and culture

The isolation and culture of EECs were described previously (***Zhao et al., 2017***). Endometrial tissues were treated with collagenase type I (Sigma, St Louis, MO, USA), hyaluronidase (Sigma), and DNase (Roche, Indianapolis, IN, USA), followed by filtration through a 40-μm cell strainer (BD Biosciences, San Jose, CA, USA) to remove the stromal cells and then through a 100-μm cell strainer (BD Biosciences) to collect the EECs. The cells were plated on matrigel-coated dishes and cultured with Keratinocyte Serum Free Medium (KSFM; Gibco, Massachusetts, USA) containing 2% fetal bovine serum (FBS), 100 U/ml penicillin and 0.1 mg/ml streptomycin.

HEK-293T and Ishikawa cells were grown in Dulbecco’s modified Eagle’s medium (DMEM) containing 10% FBS, 100 U/ml penicillin and 0.1 mg/ml streptomycin at 37°C with 5% CO_2_.

RNA and quantitative reverse transcription-PCR (qRT-PCR)

Total RNA was extracted by Trizol reagent (Invitrogen Life Technologies, Carlsbad, CA, USA). For acquisition of RNA from cytoplasm and nucleus respectively, the nuclear and cytoplasmic fractions were isolated using NE-PER Nuclear and Cytoplasmic Extraction Reagents (Thermo Scientific, Waltham, MA, USA). The circRNAs were reverse-transcribed using PrimeScriptTM reagent Kit with DNA Eraser (TaKaRa BIO, Japan). qRT-PCR was performed using SYBR qPCR Master Mix (Vazyme). U6 and GAPDH were used as internal controls. The relative RNA levels were determined by the 2^−ΔCt^ or 2^−ΔΔCt^ method. All primers are listed in ***Supplementary File 5***.

### Western blotting

After the determination of protein concentration of the cell lysates, each sample with equal amount of protein was subjected to SDS-PAGE gel (10% or 15%) and transferred to a PVDF membrane (Millipore, Massachusetts, USA). The membrane was incubated with the specific primary antibody overnight at 4°C, followed by incubation with HRP-conjugated anti-rabbit IgG (1:2000, Cell Signaling Technology, Boston, Mass, USA, cat# 7074S) or anti-mouse IgG (1:2000, Cell Signaling Technology, cat# 7076S). The signals were visualized with ECL solution (Millipore). The primary antibodies were: E-cad (1:1000, Abcam, Cambridge, UK, cat# ab76055), N-cad (1:500, Abcam, cat# ab18203), α-SMA (1:500, Abcam, cat# ab5694), FN (1:500, Abcam, cat# ab2413), ΔNp63α (1:500, Millipore, cat# ABS552). The antibody for GAPDH (1:10000, ABclonal, Mass, USA, cat# AC035) was used as a control.

### Immunohistochemistry and Immunofluorescence

For immunohistochemistry, mice endometrium tissues were fixed in 10% formaldehyde for 8 hours at 4°C, transferred to a tissue processor (Leica ASP300 S, Wetzlar, Germany) for dehydration and wax infiltration, and embedded in paraffin. Paraffin blocks were cut into 2-μm-thick slices. After dewaxing and blocking endogenous peroxidase activity in 3% H2O2, slices were heat-mediated in universal antigen retrieval. The slides were blocked in 2% bovine serum albumin/PBST for 1 hour. After these pretreatments, slides were incubated with primary antibodies at 4℃ overnight and washed by PBST. HRP-conjugated secondary antibodies were incubated for 1 hour. Slides were exposed to 3’3-diaminobenzidine (DAB) to visualize the antigen signals. The positive staining was confirmed in a blinded manner by two independent observers by microscopy (DMi8, Leica, Wetzlar, Germany). For cell immunofluorescence, isolated EECs that had been plated on matrigel-coated coverslips for varying times were fixed in 4% paraformaldehyde for 10 minutes and permeabilized with cold methanol. Fixed cells were stained with a primary antibody overnight, washed, and incubated with secondary antibodies conjugated to fluorescein or rhodamine. Nuclei of EECs were stained using 4′,6-diamidino-2-phenylindole (DAPI, Abcam). The primary antibodies were: E-cad (1:400, Abcam), N-cad (1:400, Abcam), α-SMA (1:100, Abcam), ΔNp63α (1:100, Millipore).

### RNA-scope assay

RNA-scope assay was performed on 5-µm thick tissue sections (Advanced Cell Diagnostics, San Francisco, CA, USA). Sections were hybridized with peptidylprolyl isomerase B (Cat# 710171), pre-miR-21 (Cat# 713661) or circPTPN12 (Cat# 854881) probes, respectively. Slides were viewed under a microscope (DMi8).

### Dual-luciferase reporter assay

The full-length sequence of circPTPN12 or the mutants were inserted into pGL3 luciferase vector to obtain the circPTPN12^wild^ or circPTPN12^mut^ constructs (Generay biotechnology, Shanghai, China). The mutants were those with deletion of each or all the three sites of circPTPN12 for binding miR-21-5p. 3’ UTR or the 3’ UTR with miR-21-5p binding site mutation of ΔNp63α were inserted into a pGL3 plasmid to obtain the pGL3-ΔNp63α^wild^ or pGL3-ΔNp63α^mut^ constructs (Generay). HEK-293T cells were co-transfected with circPTPN12^wild^ or circPTPN12^mut^ or pGL3-ΔNp63α^wild^ or pGL3-ΔNp63α^mut^ and miR-21-5p mimic (Ribobio, Guangzhou, China) or miR-21-5p inhibitor (Ribobio) or circPTPN12 plasmid using Lipofectamine 3000. After 24 hours, the firefly and Renilla luciferase activities were measured consecutively using a Dual Luciferase Reporter Assay kit (Vazyme).

### Recombinant ΔNp63α adenovirus (Ad-ΔNp63α) construction

ΔNp63α adenovirus were constructed as previous study (***Zhao et al., 2017***). The open reading frame (GenBank: AF075431.1) and 3’UTR of Homo sapiens of ΔNp63α were inserted into a pDC315-3FLAG-SV40-EGFP vector and ligated into a shuttle plasmid.

Then, HEK-293A cells were co-transfected with the shuttle plasmid and adenoviral backbone plasmid to produce ΔNp63α recombinant adenoviral vector. The same vector was used without the ΔNp63α insertion to construct the control virus (GeneChem, Shanghai, China). circPTPN12 adenovirus (Ad-circPTPN12), adeno-associated virus (AAV-circPTPN12) and plasmid construction circPTPN12 adenovirus was obtained from Genechem. Briefly, the full length of circPTPN12 with its flanking introns including complementary Alu elements was amplified to insert into a pCMV-MCS-EGFP vector, and an empty pCMV-MCS-EGFP vector served as control virus. circPTPN12 adeno-associated virus was obtained from HanBio Biotechnology (Shanghai, China). The full length of circPTPN12 was inserted into pHBAAV-CMV-CircRNA-EF1-ZsGreen vector and then inserted into adeno-associated virus. The empty vector was used as control virus. circPTPN12 overexpression plasmid was obtained from Genepharma (Shanghai, China). The full length of circPTPN12 with its flanking introns was inserted into a pGCMV/MCS/Neo (pEX-3) vector.

### RNA-RNA pull-down assay

A biotin-labeled circPTPN12 probe and an unlabeled probe (5’-AGGCCATTACAATGATCTGCAATGAATAC-3’, General Biosystems, Chuzhou, China) were separately incubated with streptavidin-coated magnetic beads (Thermo Scientific), followed by incubation with the lysates of HEK-293T cells transfected with circPTPN12 plasmid. The RNA was extracted from the precipitated magnetic beads, and subjected to qRT-PCR, and the resultant qRT-PCR products were further analyzed by agarose gel electrophoresis.

### RNA-binding protein immunoprecipitation (RIP)

The lysates of HEK-293T cells transfected with circPTPN12 plasmid were incubated with Protein-A/G agarose beads (Millipore) and antibody against Ago2 (Abcam, cat# ab186733). RNA was extracted from the precipitates. The abundance of circPTPN12 and miR-21-5p in the precipitates was detected by qRT-PCR and the resultant qRT-PCR products were further analyzed by agarose gel electrophoresis.

### RNA fluorescent in situ hybridization (FISH)

Ishikawa cell sections were hybridized with CY3 labeled circPTPN12 probe (CY3-5’-TTACAATGATCTGCAATGAATA-3’, Geneseed, Guangzhou, China) and FITC labeled miR-21-5p probe (FITC-5’-TCAACATCAGTCTGATAAGCTA-3’, Geneseed) at 37°C for 18 hours. Nuclei were stained using DAPI. The images were acquired with confocal microscopy (TCS SP2 AOBS, Leica).

### Mouse models

All mouse procedures were approved by the Institutional Animal Care and Use Committee at the Nanjing Drum Tower Hospital. BALB/c female mice, at 8-10 weeks of age, 18-20 g, were bred in the Animal Laboratory Center of Nanjing Drum Tower Hospital. Mechanical injury of endometrium was performed as previously reported (***Zhao et al., 2020***). Briefly, mice at estrum defined by vaginal smears were anesthetized with isoflurane. Laparotomy was performed to expose the uterus, followed by insertion of rough surfaced needle to injure uterine endometrium. Mice in sham-operation were just subjected to laparotomy to expose the uterus without injury. AAV-circPTPN12 or AAV-control (10 μl of 1 × 10^9^ viral genomes/µl) was administered via intrauterine injection 5 days after mechanical injury to study the function of circPTPN12 in endometrium fibrosis. For the therapeutic effect of miR-21-5p, agomir-21-5p (Ribobio) or agomir-NC (Ribobio) (5 nmol in 10 μl) was administered every 5 days for 3 times via intrauterine injection after mechanical injury and AAV-circPTPN12 injection. Mouse model grouping was shown in ***Supplementary File 6***. Uterine tissues were collected at estrum, 24 to 26 days after first surgery. Two side uteri of each mouse were divided into three parts: 1/3 for RNA isolation, 1/3 for immunohistochemistry analysis, and 1/3 for immunofluorescence.

### Statistical Analysis

Statistical analyses were carried out by GraphPad Prism software (version 6.0). Student’s t-test was used to analyze two experimental groups if data were normally distributed and One-way ANOVA followed by a Student-Newman-Keuls multiple comparisons test were used for comparing three or more groups. Data were presented as mean ± standard deviation (SD) or means ± standard error of mean (SEM) indicated in each figure legend. *P* < 0.05 was considered statistically significant.

## Acknowledgments

We thank Honghong Yao and her team for the technical guidance of circRNA fluorescence in situ hybridization in this study. We thank Yihua Zhou for polishing up the article. This study was supported by The Strategic Priority Research Program of the Chinese Academy of Sciences (XDA16040302), National Key R&D Program of China (2018YFC1004404), National Natural Science Foundation of China (81971336, 81771526, 82071600), Excellent Youth Natural Science Foundation of Jiangsu Province (BK20170051), Jiangsu Province’s Key Provincial Talents Program (ZDRCA2016067), and Jiangsu Biobank of Clinical Resources (BM2015004).

## Competing interests

All authors declare that they have no conflict of interest.

## Additional files

Supplementary File 1. Dysregulated miRNAs in IUA patients.

Supplementary File 2. Forty upregulated and forty downregulated miRNAs with higher expression abundance.

Supplementary File 3. circRNAs with binding sites for miR-21-5p.

Supplementary File 4. Clinical information of all patients and controls.

Supplementary File 5. All primers used in this study.

Supplementary File 6. Mouse model grouping.

---

**Figure 1-figure supplement 1.**
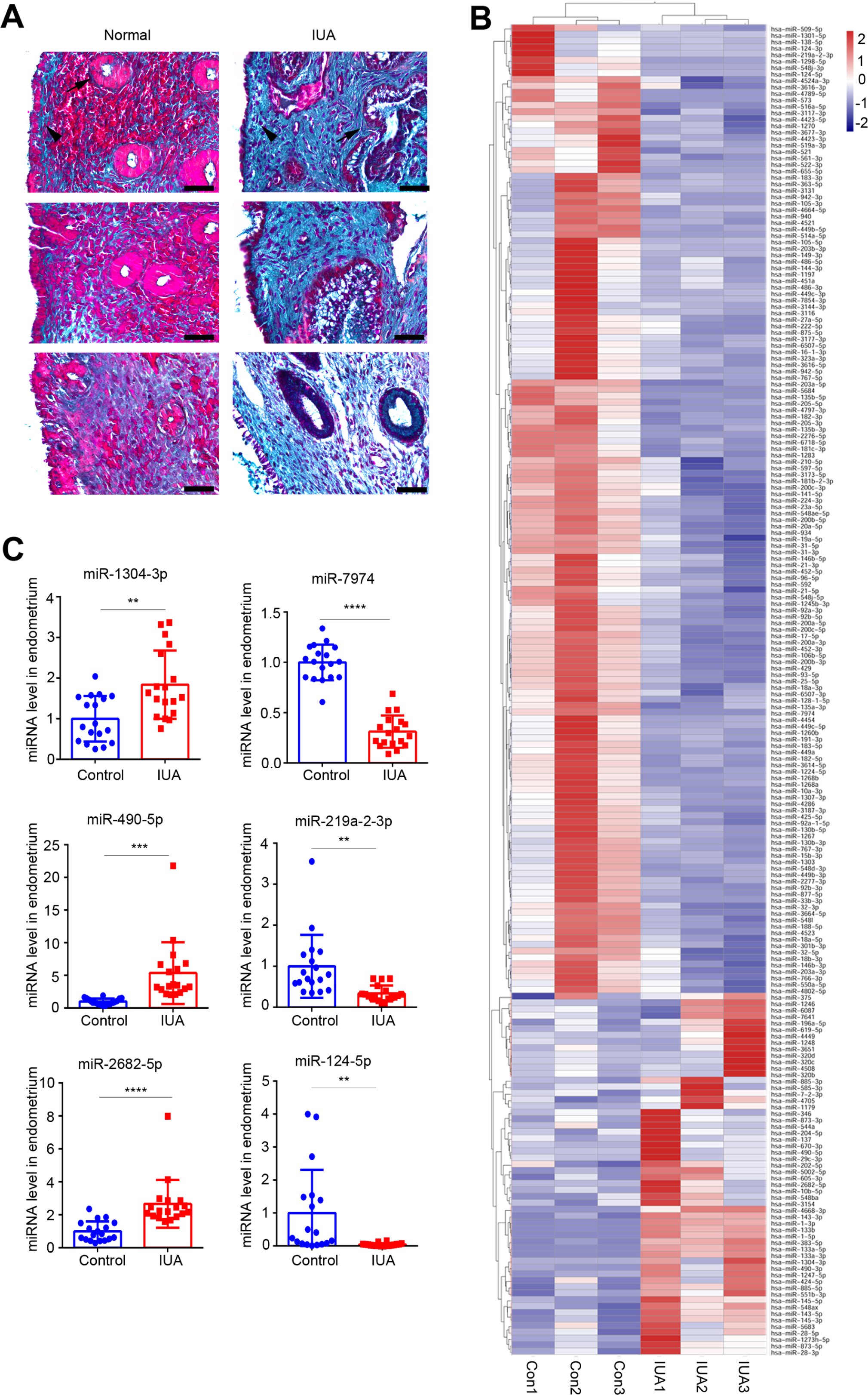
miRNAs expression profile. (**A**) Representative images of Masson staining in endometrium samples used for high-throughput sequencing from severe IUA patients (*n* =3) and controls (*n* = 3). Scale bars, 50 μm. Arrow head: luminal epithelial cells; arrow: glandular epithelial cells. (**B**) A heatmap showing all dysregulated miRNAs (56 upregulated and 150 downregulated) in endometrium samples from severe IUA patients (*n* =3) and controls (*n* = 3). (**C**) Validation of top 3 most upregulated and 3 most downregulated miRNAs in the endometria of severe IUA patients (*n* = 18) and controls (*n* = 18) by qRT-PCR. Error bars indicate mean ± SD. ** *P* < 0.01, **** *P* < 0.0001.

**Figure 1-figure supplement 2.**
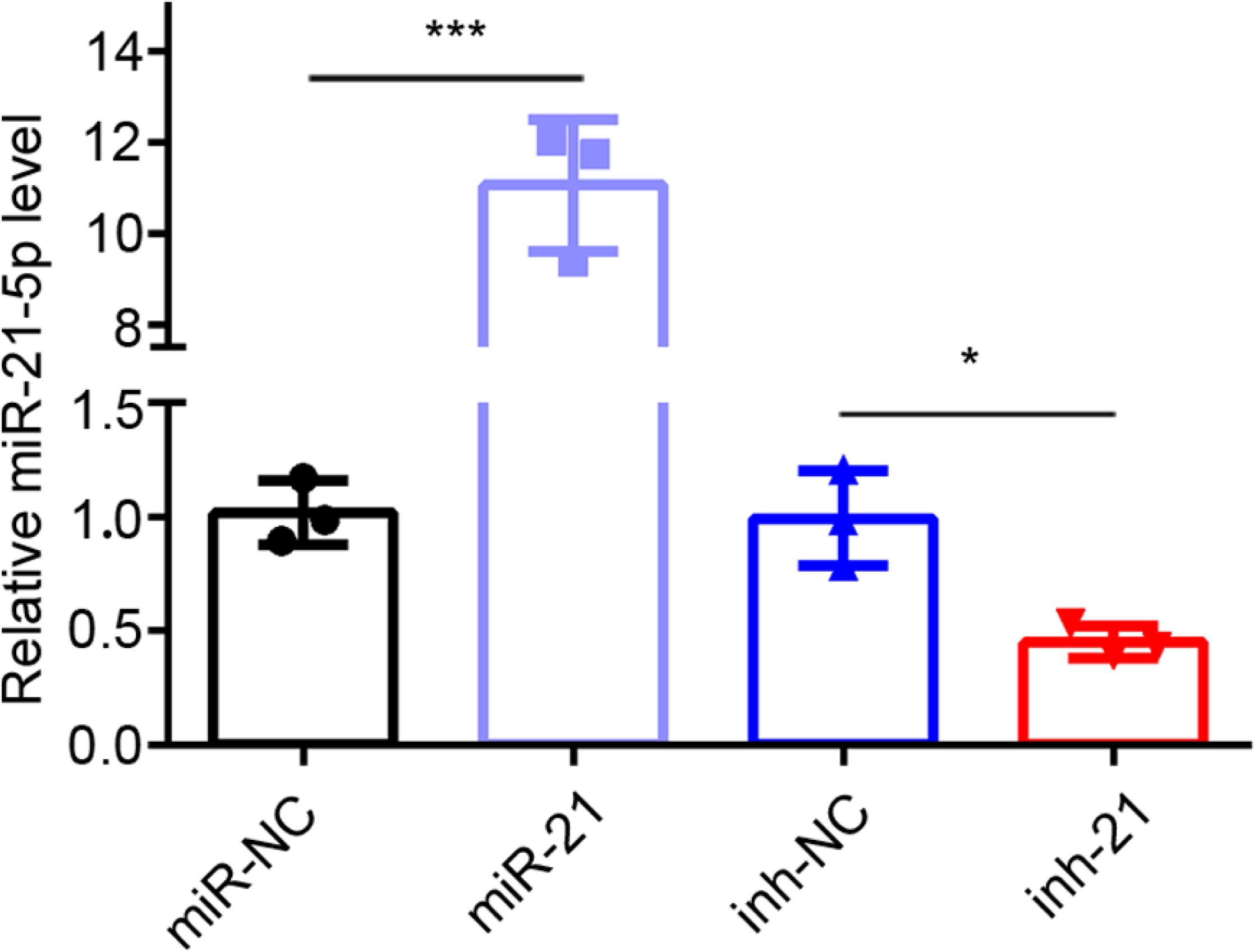
miR-21-5p transfection efficiency. qRT-PCR analysis of miR-21-5p level in EECs transfected with miR-21-5p mimic (*n*=3) or inhibitor (*n*=3) for 48 hours. Error bars indicate mean ± SD. ** *P* < 0.01, **** *P* < 0.0001. **Supplementary File 1.** Dysregulated miRNAs in IUA patients. **Supplementary File 2.** Forty upregulated and forty downregulated miRNAs with higher expression abundance.

**Figure 2-figure supplement 1.**
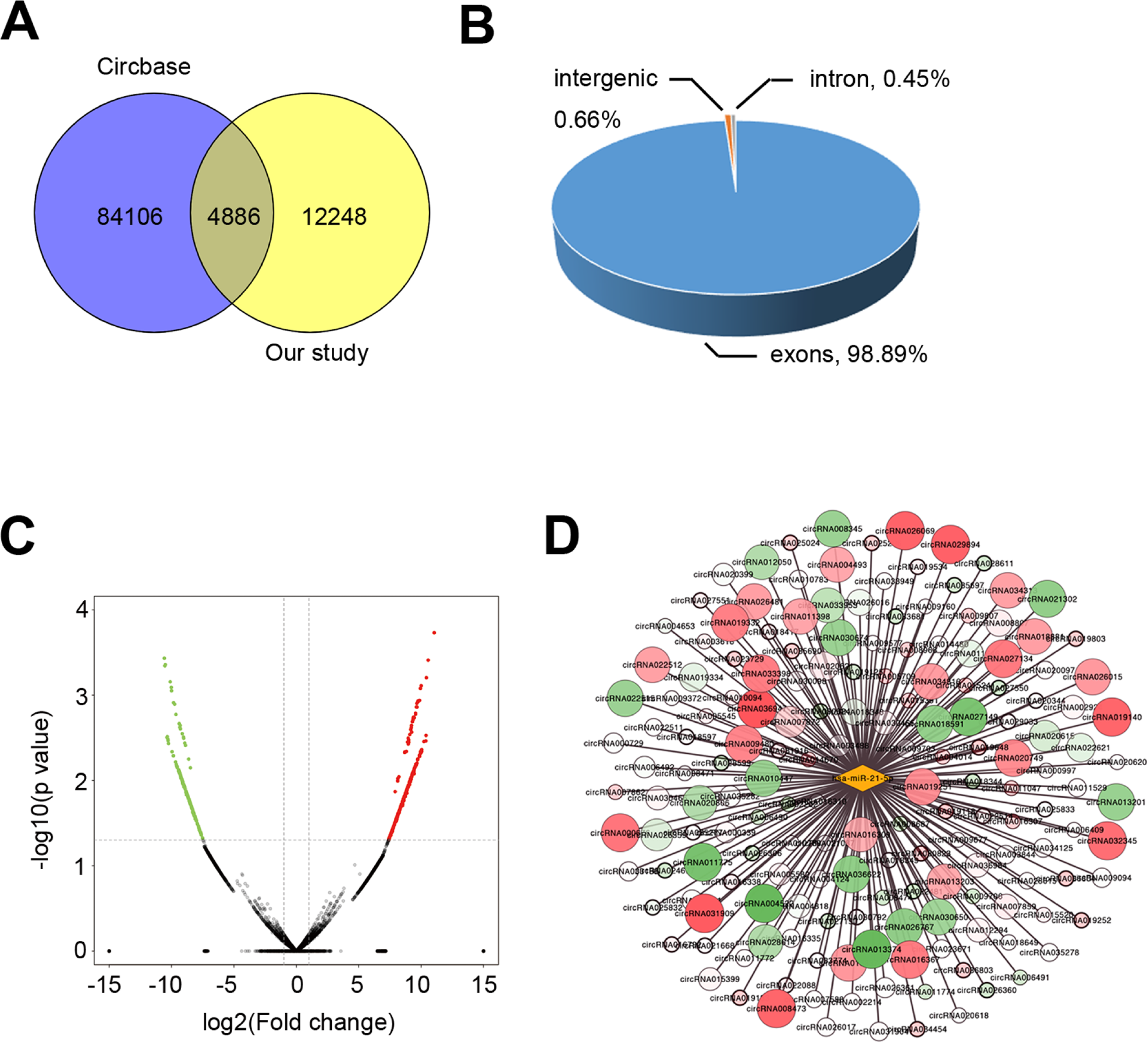
circRNAs expression profile. (**A**) Venn diagram showing the intersection of the detected circRNAs in the endometria from severe IUA patients (*n* = 3) and controls (*n* = 3) based on the high-throughput RNA-seq analysis and the circRNAs already registered in the CircBase database. (**B**) Pie chart showing the genomic origin of the circRNAs which are detected in the endometria and have registered in the CircBase database. (**C**) Volcano plots of circRNA-seq of the endometria from severe IUA patients (*n* = 3) and controls (*n* = 3) based on the high-throughput RNA-sequencing analysis. The red dots represent the circRNAs up-regulated and the green ones represent the down-regulated (P <0.05 and fold change > 2). (**D**) Network diagram of detected circRNAs possessing binding sites for miR-21-5p. **Supplementary File 3.** circRNAs with binding sites for miR-21-5p.

**Figure 3-figure supplement 1.**
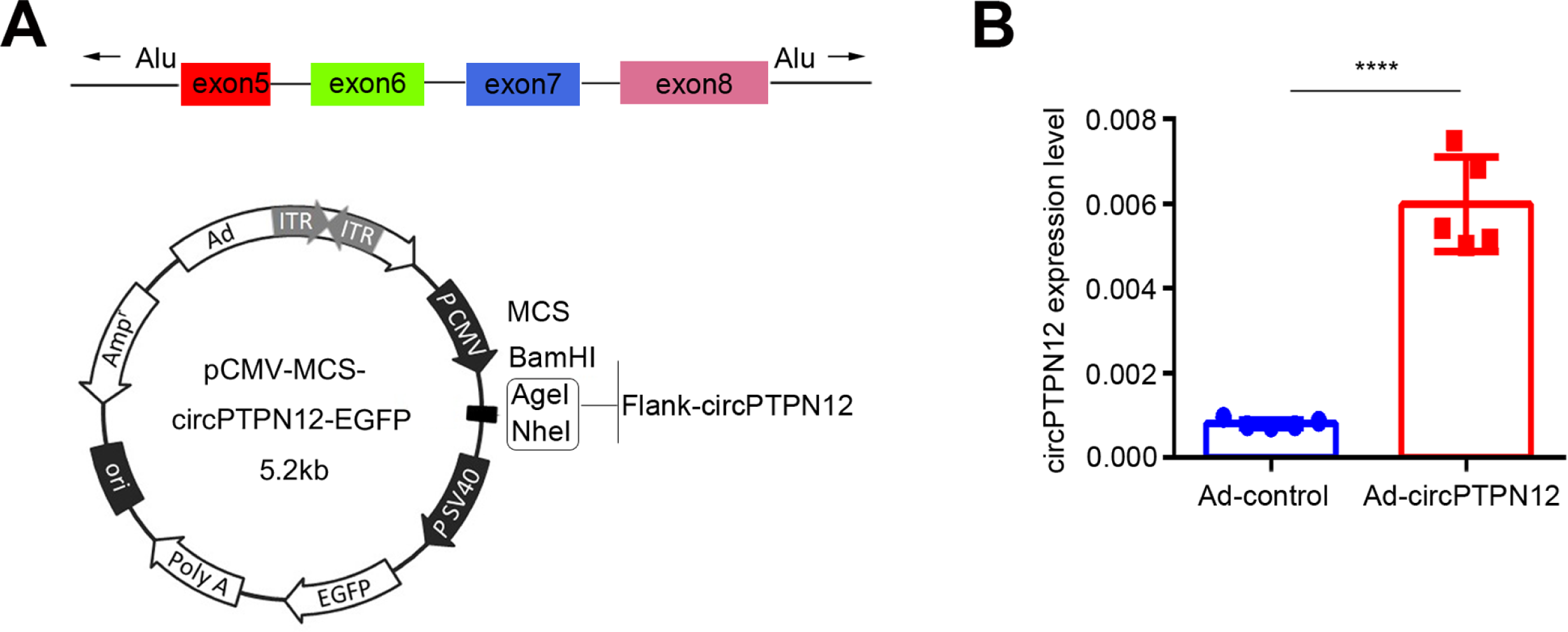
circPTPN12 adenovirus (Ad-circPTPN12) construction. (**A**) Structure of Ad-circPTPN12. (**B**) qRT-PCR analysis of circPTPN12 level in EECs infected by Ad-circPTPN12 or Ad-control for 24 hours (*n* = 5). Error bars indicate mean ± SD. **** *P* < 0.0001.

**Figure 5-figure supplement 1.**
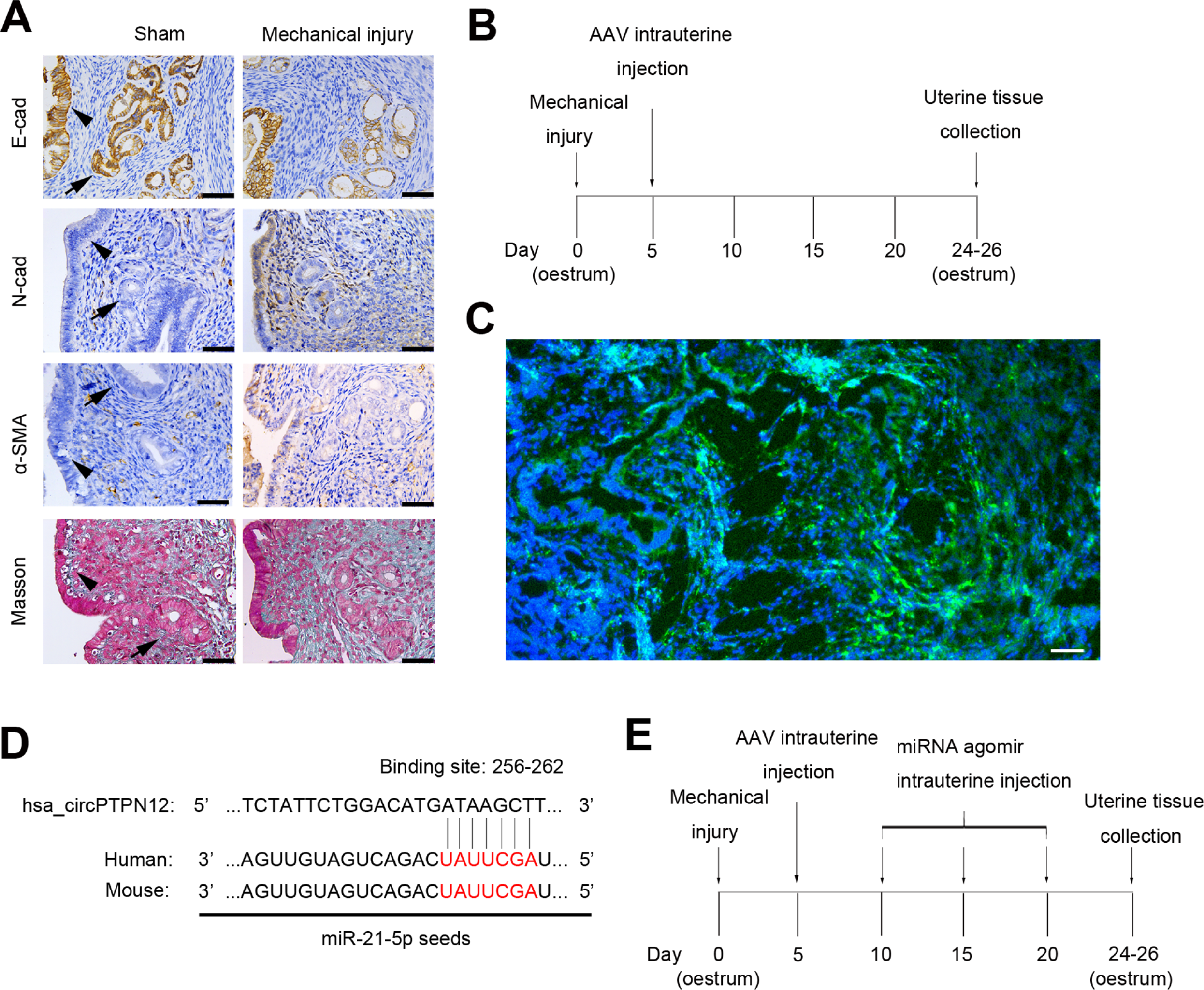
IUA-like mouse model construction. (**A**) Representative images of E-cad, N-cad, α-SMA and Masson staining in endometrial biopsies of sham-operation (*n* = 3) and mechanical injury model mice (*n* = 3). Scale bars, 50 μm. Arrow head: luminal epithelial cells; arrow: glandular epithelial cells. (**B**) A flow diagram of experimental procedure with sham-operation, mechanical injury and AAV-control or AAV-circPTPN12 uterine injection. (**C**) zsGREEN Fluorescence in endometrial biopsies of mice 21 days after AAV-circPTPN12 injection. Scale bars, 20 μm. (**D**) Conserved miR-21-5p sequence in human and mouse. (**E**) A flow diagram of experimental procedure with sham-operation, mechanical injury with AAV-circPTPN12 and agomir-21-5p or agomir-NC uterine injection.

## Notes

### Competing Interest Statement

The authors have declared no competing interest.

## References

Brown Brian D. & Naldini Luigi. 2009. Exploiting and antagonizing microRNA regulation for therapeutic and experimental applications. Nat Rev Genet, 10(8), 578–85. doi:10.1038/nrg2628.

Candi E., Amelio I., Agostini M. & Melino G. 2015. MicroRNAs and p63 in epithelial stemness. Cell Death Differ, 22(1), 12–21. doi:10.1038/cdd.2014.113.

Chau B Nelson., Xin Cuiyan., Hartner Jochen., Ren Shuyu., Castano Ana P., Linn Geoffrey., Li Jian., Tran Phong T., Kaimal Vivek., Huang Xinqiang., Chang Aaron N., Li Shenyang., Kalra Aarti., Grafals Monica., Portilla Didier., MacKenna Deidre A., Orkin Stuart H. & Duffield Jeremy S. 2012. MicroRNA-21 promotes fibrosis of the kidney by silencing metabolic pathways. Sci Transl Med, 4(121), 121ra18. doi:10.1126/scitranslmed.3003205.

Cheng Zhuoan., Yu Chengtao., Cui Shaohua., Wang Hui., Jin Haojie., Wang Cun., Li Botai., Qin Meilin., Yang Chen., He Jia., Zuo Qiaozhu., Wang Siying., Liu Jun., Ye Weidong., Lv Yuanyuan., Zhao Fangyu., Yao Ming., Jiang Liyan. & Qin Wenxin. 2019. circTP63 functions as a ceRNA to promote lung squamous cell carcinoma progression by upregulating FOXM1. Nat Commun, 10(1), 3200. doi:10.1038/s41467-019-11162-4.

Das Sudipta., Kumar Manish., Negi Vinny., Pattnaik Bijay., Prakash Y S., Agrawal Anurag. & Ghosh Balaram. 2014. MicroRNA-326 regulates profibrotic functions of transforming growth factor-β in pulmonary fibrosis. Am J Respir Cell Mol Biol, 50(5), 882–92. doi:10.1165/rcmb.2013-0195OC.

Fu X & Liu R. paper presented at the 6th International Conference on Bioinformatics and Computational Biology, Las Vegas, NV, 24 to 26 March 2014.

Gebert Luca F R. & MacRae Ian J. 2019. Regulation of microRNA function in animals. Nat Rev Mol Cell Biol, 20(1), 21–37. doi:10.1038/s41580-018-0045-7.

Iwano Masayuki., Plieth David., Danoff Theodore M., Xue Chengsen., Okada Hirokazu. & Neilson Eric G. 2002. Evidence that fibroblasts derive from epithelium during tissue fibrosis. J Clin Invest, 110(3), 341–50. doi:10.1172/JCI15518.

Lena A M., Shalom-Feuerstein R., Rivetti di Val Cervo P., Aberdam D., Knight R A., Melino G. & Candi E. 2008. miR-203 represses ’stemness’ by repressing DeltaNp63. Cell Death Differ, 15(7), 1187–95. doi:10.1038/cdd.2008.69.

Li Jingxiong., Du Shaohua., Sheng Xiujie., Liu Juan., Cen Bohong., Huang Feng. & He Yuanli. 2016. MicroRNA-29b Inhibits Endometrial Fibrosis by Regulating the Sp1-TGF-β1/Smad-CTGF Axis in a Rat Model. Reprod Sci, 23(3), 386–94. doi:10.1177/1933719115602768.

Li Xiang., Yang Li. & Chen Ling-Ling. 2018. The Biogenesis, Functions, and Challenges of Circular RNAs. Mol Cell, 71(3), 428–442. doi:10.1016/j.molcel.2018.06.034.

Liu Yong., Bi Xianjin., Xiong Jiachuan., Han Wenhao., Xiao Tangli., Xu Xinli., Yang Ke., Liu Chi., Jiang Wei., He Ting., Yu Yanlin., Li Yan., Zhang Jingbo., Zhang Bo. & Zhao Jinghong. 2019. MicroRNA-34a Promotes Renal Fibrosis by Downregulation of Klotho in Tubular Epithelial Cells. Mol Ther, 27(5), 1051–1065. doi:10.1016/j.ymthe.2019.02.009.

Liu Yuwei., Xue Mengzhu., Du Shaowei., Feng Wanwan., Zhang Ke., Zhang Liwen., Liu Haiyue., Jia Guoyi., Wu Lingshuang., Hu Xin., Chen Luonan. & Wang Peng. 2019. Competitive endogenous RNA is an intrinsic component of EMT regulatory circuits and modulates EMT. Nat Commun, 10(1), 1637. doi:10.1038/s41467-019-09649-1.

March Charles M. 2011a. Management of Asherman’s syndrome. Reprod Biomed Online, 23(1), 63–76. doi:10.1016/j.rbmo.2010.11.018.

March Charles M. 2011b. Asherman’s syndrome. Semin Reprod Med, 29(2), 83–94. doi:10.1055/s-0031-1272470.

Nieto M Angela., Huang Ruby Yun-Ju., Jackson Rebecca A. & Thiery Jean Paul. 2016. EMT: 2016. Cell, 166(1), 21–45. doi:10.1016/j.cell.2016.06.028.

Ning Jing., Zhang Hongtao. & Yang Hongwei. 2018. MicroRNA-326 inhibits endometrial fibrosis by regulating TGF-β1/Smad3 pathway in intrauterine adhesions. Mol Med Rep, 18(2), 2286–2292. doi:10.3892/mmr.2018.9187.

Pandit Kusum V., Corcoran David., Yousef Hanadie., Yarlagadda Manohar., Tzouvelekis Argyris., Gibson Kevin F., Konishi Kazuhisa., Yousem Samuel A., Singh Mandal., Handley Daniel., Richards Thomas., Selman Moises., Watkins Simon C., Pardo Annie., Ben-Yehudah Ahmi., Bouros Demosthenes., Eickelberg Oliver., Ray Prabir., Benos Panayiotis V. & Kaminski Naftali. 2010. Inhibition and role of let-7d in idiopathic pulmonary fibrosis. Am J Respir Crit Care Med, 182(2), 220–9. doi:10.1164/rccm.200911-1698OC.

Rodriguez Calleja Lidia., Jacques Camille., Lamoureux François., Baud’huin Marc., Tellez Gabriel Marta., Quillard Thibaut., Sahay Debashish., Perrot Pierre., Amiaud Jerome., Charrier Celine., Brion Regis., Lecanda Fernando., Verrecchia Franck., Heymann Dominique., Ellisen Leif W. & Ory Benjamin. 2016. ΔNp63α Silences a miRNA Program to Aberrantly Initiate a Wound-Healing Program That Promotes TGFβ-Induced Metastasis. Cancer Res, 76(11), 3236–51. doi:10.1158/0008-5472.CAN-15-2317.

Sun Jingping., Li Qiuyue., Lian Ximeng., Zhu Zhonghui., Chen Xiaowei., Pei Wanying., Li Siling., Abbas Ali., Wang Yan. & Tian Lin. 2019. MicroRNA-29b Mediates Lung Mesenchymal-Epithelial Transition and Prevents Lung Fibrosis in the Silicosis Model. Mol Ther Nucleic Acids, 14, 20–31. doi:10.1016/j.omtn.2018.10.017.

The American Fertility Society classifications of adnexal adhesions, distal tubal occlusion, tubal occlusion secondary to tubal ligation, tubal pregnancies, müllerian anomalies and intrauterine adhesions. 1988. Fertil Steril, 49(6), 944-55.doi: 10.101610.1016/s0015-0282(16)59942-7.

Wang Yuanyuan., Liu Lingling., Peng Wei., Liu Huiming., Liang Luqun., Zhang Xiaohuan., Mao Yanwen., Zhou Xingcheng., Shi Mingjun., Xiao Ying., Zhang Fan., Zhang Yingying., Liu Lirong., Yan Rui. & Guo Bing. 2019. Ski-related novel protein suppresses the development of diabetic nephropathy by modulating transforming growth factor-β signaling and microRNA-21 expression. J Cell Physiol, 234(10), 17925–17936. doi:10.1002/jcp.28425.

Wu Jing., Liu Shang., Yuan Zhi-Wei., Liu Jian-He., Li Jiong-Ming., Chen Tao. & Fang Ke-Wei. 2018. MicroRNA-27a Suppresses Detrusor Fibrosis in Streptozotocin-Induced Diabetic Rats by Targeting PRKAA2 Through the TGF-β1/Smad3 Signaling Pathway. Cell Physiol Biochem, 45(4), 1333–1349. doi:10.1159/000487560.

Yamada Mitsuhiro., Kubo Hiroshi., Ota Chiharu., Takahashi Toru., Tando Yukiko., Suzuki Takaya., Fujino Naoya., Makiguchi Tomonori., Takagi Kiyoshi., Suzuki Takashi. & Ichinose Masakazu. 2013. The increase of microRNA-21 during lung fibrosis and its contribution to epithelial-mesenchymal transition in pulmonary epithelial cells. Respir Res, 14, 95. doi:10.1186/1465-9921-14-95.

Yang Huan., Wu Su., Feng Ran., Huang Junjiu., Liu Lixiang., Liu Feng. & Chen Yuqing. 2017. Vitamin C plus hydrogel facilitates bone marrow stromal cell-mediated endometrium regeneration in rats. Stem Cell Res Ther, 8(1), 267. doi:10.1186/s13287-017-0718-8.

Yao Yuan., Chen Ran., Wang Guowu., Zhang Yu. & Liu Fang. 2019. Exosomes derived from mesenchymal stem cells reverse EMT via TGF-β1/Smad pathway and promote repair of damaged endometrium. Stem Cell Res Ther, 10(1), 225. doi:10.1186/s13287-019-1332-8.

Yu Dan., Wong Yat-May., Cheong Ying., Xia Enlan. & Li Tin-Chiu. 2008. Asherman syndrome--one century later. Fertil Steril, 89(4), 759–79. doi:10.1016/j.fertnstert.2008.02.096.

Zeisberg Michael., Yang Changqing., Martino Margot., Duncan Michael B., Rieder Florian., Tanjore Harikrishna. & Kalluri Raghu. 2007. Fibroblasts derive from hepatocytes in liver fibrosis via epithelial to mesenchymal transition. J Biol Chem, 282(32), 23337–47. doi:10.1074/jbc.M700194200.

Zhang Aiqing., Li Min., Wang Bin., Klein Janet D., Price S Russ. & Wang Xiaonan H. 2018. miRNA-23a/27a attenuates muscle atrophy and renal fibrosis through muscle-kidney crosstalk. J Cachexia Sarcopenia Muscle, 9(4), 755–770. doi:10.1002/jcsm.12296.

Zhao Guangfeng., Cao Yun., Zhu Xianghong., Tang Xiaoqiu., Ding Lijun., Sun Haixiang., Li Juan., Li Xinan., Dai Chenyan., Ru Tong., Zhu Hui., Lu Jingjie., Lin Caimei., Wang Jingmei., Yan Guijun., Wang Huiyan., Wang Lei., Dai Yimin., Wang Bin., Li Ruotian., Dai Jianwu., Zhou Yan. & Hu Yali. 2017. Transplantation of collagen scaffold with autologous bone marrow mononuclear cells promotes functional endometrium reconstruction via downregulating ΔNp63 expression in Asherman’s syndrome. Sci China Life Sci, 60(4), 404–416. doi:10.1007/s11427-016-0328-y.

Zhao Guangfeng., Li Ruotian., Cao Yun., Song Minmin., Jiang Peipei., Wu Qianwen., Zhou Zhenhua., Zhu Hui., Wang Huiyan., Dai Chenyan., Liu Dan., Yao Simin., Lv Haining., Wang Limin., Dai Jianwu., Zhou Yan. & Hu Yali. 2020. ΔNp63α-induced DUSP4/GSK3β/SNAI1 pathway in epithelial cells drives endometrial fibrosis. Cell Death Dis, 11(6), 449. doi:10.1038/s41419-020-2666-y.

Zheng Qiupeng., Bao Chunyang., Guo Weijie., Li Shuyi., Chen Jie., Chen Bing., Luo Yanting., Lyu Dongbin., Li Yan., Shi Guohai., Liang Linhui., Gu Jianren., He Xianghuo. & Huang Shenglin. 2016. Circular RNA profiling reveals an abundant circHIPK3 that regulates cell growth by sponging multiple miRNAs. Nat Commun, 7, 11215. doi:10.1038/ncomms11215.

Zhuang Ya-Se., Liao Ying-Ying., Liu Bo-Yi., Fang Zhi-Cheng., Chen Li., Min Li. & Chen Wei. 2018. MicroRNA-27a mediates the Wnt/β-catenin pathway to affect the myocardial fibrosis in rats with chronic heart failure. Cardiovasc Ther, e12468. doi:10.1111/1755-5922.12468.

