## Supplementary File 1 for "Targeting circPTPN12/miR-21-5p/ΔNp63α pathway as a therapeutic strategy for human endometrial fibrosis"

Supplementary File 1. Dysregulated miRNAs in IUA patients.

| GeneName | Con_TPM | IUA_TPM | log2(IUA/Con) | up-or-down | p_value | q_value |
| --- | --- | --- | --- | --- | --- | --- |
| hsa-miR-1304-3p | 0.001 | 0.383333 | 8.582456 | up | 0.016011 | 0.163118 |
| hsa-miR-490-5p | 0.001 | 0.326667 | 8.351675 | up | 0.018602 | 0.181255 |
| hsa-miR-2682-5p | 0.001 | 0.236667 | 7.886713 | up | 0.03182 | 0.264797 |
| hsa-miR-4668-3p | 0.001 | 0.203333 | 7.667703 | up | 0.047149 | 0.324826 |
| hsa-miR-133a-5p | 0.266667 | 30.26333 | 6.82639 | up | 4.75E-26 | 2.00E-23 |
| hsa-miR-133b | 0.893333 | 92.96 | 6.701268 | up | 5.48E-31 | 3.07E-28 |
| hsa-miR-1-3p | 25.51333 | 2035.3 | 6.317846 | up | 1.76E-38 | 2.96E-35 |
| hsa-miR-133a-3p | 14.65667 | 1150.85 | 6.294999 | up | 6.73E-38 | 5.66E-35 |
| hsa-miR-1-5p | 0.083333 | 3.833333 | 5.523562 | up | 7.34E-14 | 2.47E-11 |
| hsa-miR-383-5p | 0.2 | 7.873333 | 5.298903 | up | 5.59E-13 | 1.57E-10 |
| hsa-miR-5002-5p | 0.02 | 0.496667 | 4.634206 | up | 0.000597 | 0.014539 |
| hsa-miR-670-3p | 0.03 | 0.626667 | 4.384664 | up | 0.001419 | 0.028071 |
| hsa-miR-1273h-5p | 0.02 | 0.313333 | 3.969626 | up | 0.034796 | 0.281211 |
| hsa-miR-490-3p | 0.563333 | 5.17 | 3.198104 | up | 4.47E-07 | 4.18E-05 |
| hsa-miR-4508 | 0.093333 | 0.756667 | 3.019194 | up | 0.034444 | 0.279715 |
| hsa-miR-548ax | 0.08 | 0.626667 | 2.969626 | up | 0.002133 | 0.037509 |
| hsa-miR-4449 | 0.49 | 3.793333 | 2.952612 | up | 0.004147 | 0.059314 |
| hsa-miR-548ba | 0.136667 | 0.966667 | 2.822357 | up | 0.002762 | 0.044645 |
| hsa-miR-4705 | 0.12 | 0.8 | 2.736966 | up | 0.008788 | 0.104767 |
| hsa-miR-1246 | 0.163333 | 1.076667 | 2.720681 | up | 0.018654 | 0.181255 |
| hsa-miR-196a-5p | 7.746667 | 46.44 | 2.58372 | up | 3.92E-05 | 0.001829 |
| hsa-miR-1248 | 6.536667 | 38.54 | 2.55973 | up | 0.002011 | 0.036345 |
| hsa-miR-885-3p | 0.183333 | 1.043333 | 2.508659 | up | 0.002172 | 0.037509 |
| hsa-miR-885-5p | 8.08 | 45.11667 | 2.481233 | up | 6.78E-08 | 7.60E-06 |
| hsa-miR-605-3p | 0.406667 | 2.053333 | 2.336049 | up | 0.00014 | 0.005008 |
| hsa-miR-143-3p | 25185.46 | 116029.7 | 2.203831 | up | 1.03E-12 | 2.47E-10 |
| hsa-miR-145-5p | 2105.527 | 9658.62 | 2.197636 | up | 1.23E-10 | 2.58E-08 |
| hsa-miR-619-5p | 0.436667 | 1.853333 | 2.085518 | up | 0.008945 | 0.105151 |
| hsa-miR-320d | 10.66 | 43.35333 | 2.023935 | up | 0.005635 | 0.074592 |
| hsa-miR-320c | 20.43667 | 83.01333 | 2.022183 | up | 0.004203 | 0.059379 |
| hsa-miR-3651 | 0.376667 | 1.48 | 1.974237 | up | 0.044453 | 0.311356 |
| hsa-miR-551b-3p | 0.406667 | 1.586667 | 1.96408 | up | 0.002933 | 0.046512 |
| hsa-miR-7-2-3p | 0.253333 | 0.943333 | 1.896731 | up | 0.039702 | 0.304609 |
| hsa-miR-137 | 9.03 | 30.15667 | 1.739679 | up | 0.012557 | 0.131927 |
| hsa-miR-6087 | 0.886667 | 2.916667 | 1.717857 | up | 0.049997 | 0.338891 |
| hsa-miR-1179 | 1.426667 | 4.62 | 1.695245 | up | 0.004776 | 0.064741 |
| hsa-miR-873-3p | 0.973333 | 3.15 | 1.694346 | up | 0.001894 | 0.034994 |
| hsa-miR-143-5p | 511.2267 | 1639.493 | 1.681215 | up | 8.83E-07 | 7.81E-05 |
| hsa-miR-5683 | 0.47 | 1.443333 | 1.618672 | up | 0.013343 | 0.138458 |
| hsa-miR-145-3p | 702.35 | 2125.387 | 1.597463 | up | 1.92E-06 | 0.000161 |
| hsa-miR-7641 | 21.24667 | 64.08667 | 1.592788 | up | 0.03977 | 0.304609 |
| hsa-miR-202-5p | 1.033333 | 3.046667 | 1.559926 | up | 0.01196 | 0.128879 |
| hsa-miR-873-5p | 11.28333 | 32.51333 | 1.526838 | up | 3.27E-05 | 0.001617 |
| hsa-miR-585-3p | 1.24 | 3.456667 | 1.479041 | up | 0.043589 | 0.308277 |
| hsa-miR-1247-5p | 32.18 | 85.14667 | 1.403786 | up | 0.000654 | 0.015712 |
| hsa-miR-3154 | 0.596667 | 1.406667 | 1.237283 | up | 0.040035 | 0.304609 |
| hsa-miR-424-5p | 10544.78 | 24695.72 | 1.227731 | up | 0.000569 | 0.014405 |
| hsa-miR-320b | 80.28333 | 185.6567 | 1.209465 | up | 0.016744 | 0.169558 |
| hsa-miR-204-5p | 168.9967 | 388.7533 | 1.20186 | up | 0.009568 | 0.109956 |
| hsa-miR-28-3p | 225.3833 | 509.0467 | 1.175417 | up | 0.000301 | 0.008287 |
| hsa-miR-10b-5p | 16433.64 | 36829.03 | 1.164191 | up | 0.000109 | 0.004279 |
| hsa-miR-544a | 0.69 | 1.543333 | 1.161381 | up | 0.043647 | 0.308277 |
| hsa-miR-28-5p | 321.2733 | 717.3133 | 1.158802 | up | 0.000736 | 0.016712 |
| hsa-miR-346 | 1.886667 | 4.203333 | 1.155694 | up | 0.028723 | 0.247388 |
| hsa-miR-29c-3p | 758.97 | 1627.207 | 1.100283 | up | 0.004332 | 0.06051 |
| hsa-miR-375 | 272.08 | 566.67 | 1.058478 | up | 0.028423 | 0.246282 |
| hsa-miR-7974 | 1.586666667 | 0.001 | -10.6318 | down | 1.40E-09 | 2.45E-07 |
| hsa-miR-219a-2-3p | 1.17 | 0.001 | -10.1923 | down | 0.000364 | 0.009569 |
| hsa-miR-124-5p | 0.87 | 0.001 | -9.76487 | down | 0.000119 | 0.004547 |
| hsa-miR-149-3p | 0.536666667 | 0.001 | -9.06788 | down | 0.001194 | 0.025096 |
| hsa-miR-1301-5p | 0.296666667 | 0.001 | -8.2127 | down | 0.043543 | 0.308277 |
| hsa-miR-514a-5p | 0.25 | 0.001 | -7.96578 | down | 0.020093 | 0.186611 |
| hsa-miR-4797-3p | 0.24 | 0.001 | -7.90689 | down | 0.013626 | 0.140524 |
| hsa-miR-548j-3p | 0.236666667 | 0.001 | -7.88671 | down | 0.041132 | 0.305366 |
| hsa-miR-655-5p | 0.216666667 | 0.001 | -7.75933 | down | 0.039505 | 0.304609 |
| hsa-miR-4789-5p | 0.21 | 0.001 | -7.71425 | down | 0.032404 | 0.266972 |
| hsa-miR-573 | 0.21 | 0.001 | -7.71425 | down | 0.032558 | 0.266972 |
| hsa-miR-7854-3p | 0.21 | 0.001 | -7.71425 | down | 0.041769 | 0.306612 |
| hsa-miR-5684 | 0.2 | 0.001 | -7.64386 | down | 0.043166 | 0.308277 |
| hsa-miR-2276-5p | 0.183333333 | 0.001 | -7.51833 | down | 0.044287 | 0.311356 |
| hsa-miR-3131 | 4.64 | 0.043333333 | -6.7425 | down | 1.73E-05 | 0.00104 |
| hsa-miR-1298-5p | 3.25 | 0.163333333 | -4.31455 | down | 6.61E-08 | 7.60E-06 |
| hsa-miR-124-3p | 50.46 | 2.773333333 | -4.18545 | down | 1.11E-05 | 0.000718 |
| hsa-miR-449c-3p | 20.86 | 1.22 | -4.09579 | down | 3.82E-07 | 3.77E-05 |
| hsa-miR-449b-3p | 11.34333333 | 0.746666667 | -3.92524 | down | 3.05E-09 | 4.66E-07 |
| hsa-miR-3664-5p | 0.386666667 | 0.026666667 | -3.85798 | down | 0.011047 | 0.122175 |
| hsa-miR-203a-5p | 0.64 | 0.046666667 | -3.77761 | down | 0.001522 | 0.029411 |
| hsa-miR-4523 | 0.36 | 0.026666667 | -3.75489 | down | 0.019651 | 0.18464 |
| hsa-miR-1224-5p | 2.426666667 | 0.193333333 | -3.64981 | down | 3.24E-07 | 3.41E-05 |
| hsa-miR-138-5p | 6.04 | 0.493333333 | -3.61391 | down | 0.000131 | 0.004911 |
| hsa-miR-1283 | 0.87 | 0.083333333 | -3.38405 | down | 0.000474 | 0.01227 |
| hsa-miR-105-3p | 0.516666667 | 0.05 | -3.36923 | down | 0.019287 | 0.183262 |
| hsa-miR-6718-5p | 0.373333333 | 0.043333333 | -3.10692 | down | 0.012425 | 0.131366 |
| hsa-miR-942-3p | 0.826666667 | 0.1 | -3.04731 | down | 0.010865 | 0.12099 |
| hsa-miR-934 | 3.586666667 | 0.44 | -3.02707 | down | 1.09E-05 | 0.000718 |
| hsa-miR-522-3p | 1.133333333 | 0.14 | -3.01707 | down | 0.002187 | 0.037509 |
| hsa-miR-135b-3p | 0.896666667 | 0.12 | -2.90154 | down | 0.001276 | 0.026152 |
| hsa-miR-3614-5p | 1.763333333 | 0.24 | -2.8772 | down | 3.81E-05 | 0.001828 |
| hsa-miR-3116 | 0.51 | 0.07 | -2.86507 | down | 0.008428 | 0.101197 |
| hsa-miR-203b-3p | 1.206666667 | 0.166666667 | -2.85599 | down | 0.00205 | 0.036658 |
| hsa-miR-519a-3p | 2.006666667 | 0.283333333 | -2.82423 | down | 0.001503 | 0.029373 |
| hsa-miR-449a | 1599.376667 | 242.3266667 | -2.72248 | down | 1.46E-09 | 2.45E-07 |
| hsa-miR-449c-5p | 414.4966667 | 64.27333333 | -2.68907 | down | 3.76E-08 | 4.86E-06 |
| hsa-miR-3144-3p | 0.753333333 | 0.12 | -2.65025 | down | 0.010662 | 0.120289 |
| hsa-miR-205-3p | 0.55 | 0.09 | -2.61143 | down | 0.007863 | 0.09578 |
| hsa-miR-205-5p | 76.63666667 | 13.00666667 | -2.55878 | down | 3.17E-05 | 0.001614 |
| hsa-miR-33b-3p | 4.03 | 0.696666667 | -2.53224 | down | 9.86E-06 | 0.000691 |
| hsa-miR-1245b-3p | 0.516666667 | 0.093333333 | -2.46877 | down | 0.01299 | 0.135625 |
| hsa-miR-875-5p | 1.96 | 0.356666667 | -2.45821 | down | 0.006419 | 0.083157 |
| hsa-miR-521 | 0.486666667 | 0.09 | -2.43494 | down | 0.039892 | 0.304609 |
| hsa-miR-135b-5p | 215.7666667 | 43.15333333 | -2.32193 | down | 5.98E-09 | 8.38E-07 |
| hsa-miR-3177-3p | 0.61 | 0.123333333 | -2.30625 | down | 0.019661 | 0.18464 |
| hsa-miR-3117-3p | 4.37 | 0.893333333 | -2.29036 | down | 0.004062 | 0.059314 |
| hsa-miR-4664-5p | 0.86 | 0.176666667 | -2.28331 | down | 0.039659 | 0.304609 |
| hsa-miR-31-3p | 51.75 | 10.67333333 | -2.27755 | down | 6.65E-06 | 0.000486 |
| hsa-miR-1268a | 3.196666667 | 0.666666667 | -2.26153 | down | 5.39E-05 | 0.002321 |
| hsa-miR-3677-3p | 0.84 | 0.176666667 | -2.24936 | down | 0.019296 | 0.183262 |
| hsa-miR-363-5p | 0.73 | 0.156666667 | -2.2202 | down | 0.03178 | 0.264797 |
| hsa-miR-1267 | 0.813333333 | 0.18 | -2.17585 | down | 0.00984 | 0.11176 |
| hsa-miR-516a-5p | 4.186666667 | 0.956666667 | -2.12971 | down | 0.000364 | 0.009569 |
| hsa-miR-1268b | 3.436666667 | 0.8 | -2.10294 | down | 0.000193 | 0.006233 |
| hsa-miR-1303 | 7.056666667 | 1.643333333 | -2.10236 | down | 0.000215 | 0.006692 |
| hsa-miR-191-3p | 6.99 | 1.653333333 | -2.07991 | down | 0.000139 | 0.005008 |
| hsa-miR-1260b | 19.65666667 | 4.67 | -2.07352 | down | 2.07E-05 | 0.001132 |
| hsa-miR-105-5p | 8.01 | 1.97 | -2.02361 | down | 0.003144 | 0.048481 |
| hsa-miR-128-1-5p | 2.89 | 0.733333333 | -1.97853 | down | 0.000722 | 0.016637 |
| hsa-miR-767-3p | 1.256666667 | 0.32 | -1.97346 | down | 0.006703 | 0.084092 |
| hsa-miR-200c-5p | 7.76 | 1.99 | -1.96329 | down | 2.44E-05 | 0.001281 |
| hsa-miR-4521 | 26.41 | 6.83 | -1.95113 | down | 0.000289 | 0.008093 |
| hsa-miR-3187-3p | 0.716666667 | 0.19 | -1.9153 | down | 0.018896 | 0.181507 |
| hsa-miR-183-5p | 152.2433333 | 41.13333333 | -1.888 | down | 3.72E-06 | 0.000284 |
| hsa-miR-130b-5p | 114.7266667 | 31.20333333 | -1.87843 | down | 5.13E-05 | 0.002271 |
| hsa-miR-23a-5p | 2.533333333 | 0.69 | -1.87637 | down | 0.000755 | 0.016797 |
| hsa-miR-92a-1-5p | 11.75333333 | 3.24 | -1.859 | down | 0.000165 | 0.005684 |
| hsa-miR-200a-5p | 28.99 | 8.34 | -1.79744 | down | 3.29E-06 | 0.000263 |
| hsa-miR-548l | 0.943333333 | 0.273333333 | -1.78711 | down | 0.017826 | 0.175232 |
| hsa-miR-4423-5p | 2.573333333 | 0.753333333 | -1.77228 | down | 0.007406 | 0.090878 |
| hsa-miR-135a-3p | 4.076666667 | 1.213333333 | -1.74841 | down | 0.00652 | 0.083663 |
| hsa-miR-25-5p | 15.83333333 | 4.726666667 | -1.74407 | down | 2.09E-05 | 0.001132 |
| hsa-miR-2277-3p | 2.593333333 | 0.786666667 | -1.72098 | down | 0.003103 | 0.048294 |
| hsa-miR-18b-3p | 0.876666667 | 0.266666667 | -1.71699 | down | 0.032137 | 0.266121 |
| hsa-miR-877-5p | 10.35 | 3.213333333 | -1.68749 | down | 0.000176 | 0.00593 |
| hsa-miR-561-3p | 1.136666667 | 0.353333333 | -1.68571 | down | 0.024503 | 0.220266 |
| hsa-miR-509-5p | 1.473333333 | 0.46 | -1.67938 | down | 0.028992 | 0.247388 |
| hsa-miR-548d-3p | 1.533333333 | 0.48 | -1.67557 | down | 0.009324 | 0.10884 |
| hsa-miR-940 | 4.88 | 1.533333333 | -1.67021 | down | 0.003731 | 0.055998 |
| hsa-miR-31-5p | 405.3566667 | 128.7433333 | -1.65469 | down | 2.09E-05 | 0.001132 |
| hsa-miR-15b-3p | 69.58666667 | 22.18666667 | -1.64912 | down | 0.000232 | 0.007087 |
| hsa-miR-4454 | 24.15 | 7.713333333 | -1.6466 | down | 0.000327 | 0.008853 |
| hsa-miR-4423-3p | 4.326666667 | 1.406666667 | -1.62098 | down | 0.008905 | 0.105151 |
| hsa-miR-486-3p | 3.216666667 | 1.053333333 | -1.6106 | down | 0.027161 | 0.240305 |
| hsa-miR-144-3p | 2204.85 | 723.12 | -1.60837 | down | 0.017532 | 0.173732 |
| hsa-miR-592 | 3.12 | 1.036666667 | -1.58959 | down | 0.002836 | 0.045405 |
| hsa-miR-92b-3p | 1062.616667 | 353.7533333 | -1.58681 | down | 1.36E-05 | 0.000844 |
| hsa-miR-3616-3p | 1.51 | 0.516666667 | -1.54724 | down | 0.04316 | 0.308277 |
| hsa-miR-18a-3p | 5.85 | 2.033333333 | -1.52459 | down | 0.002145 | 0.037509 |
| hsa-miR-27a-5p | 26.56666667 | 9.253333333 | -1.52157 | down | 0.000762 | 0.016797 |
| hsa-miR-32-3p | 23.97 | 8.473333333 | -1.50023 | down | 0.000255 | 0.007397 |
| hsa-miR-4802-5p | 1.106666667 | 0.4 | -1.46815 | down | 0.035172 | 0.282892 |
| hsa-miR-449b-5p | 49.62333333 | 18.11 | -1.45423 | down | 0.003303 | 0.05048 |
| hsa-miR-200b-5p | 42.99333333 | 15.89 | -1.43599 | down | 6.32E-05 | 0.002528 |
| hsa-miR-3616-5p | 3.626666667 | 1.343333333 | -1.43283 | down | 0.007137 | 0.08821 |
| hsa-miR-548j-5p | 2.276666667 | 0.843333333 | -1.43275 | down | 0.006797 | 0.084635 |
| hsa-miR-183-3p | 3.566666667 | 1.323333333 | -1.4304 | down | 0.028258 | 0.246125 |
| hsa-miR-4524a-3p | 1.76 | 0.67 | -1.39334 | down | 0.030982 | 0.260407 |
| hsa-miR-4286 | 60.04 | 22.97666667 | -1.38575 | down | 0.000189 | 0.006233 |
| hsa-miR-451a | 15701.32 | 6017.756667 | -1.38359 | down | 0.027449 | 0.241385 |
| hsa-miR-141-5p | 25.41333333 | 9.783333333 | -1.37719 | down | 0.002362 | 0.039896 |
| hsa-miR-222-5p | 12.26333333 | 4.736666667 | -1.37241 | down | 0.008388 | 0.101197 |
| hsa-miR-6507-3p | 1.336666667 | 0.53 | -1.33458 | down | 0.040785 | 0.305049 |
| hsa-miR-429 | 809.0566667 | 321.21 | -1.33272 | down | 5.74E-05 | 0.002414 |
| hsa-miR-16-1-3p | 9.986666667 | 3.986666667 | -1.32482 | down | 0.002373 | 0.039896 |
| hsa-miR-548ae-5p | 4.913333333 | 1.966666667 | -1.32095 | down | 0.004164 | 0.059314 |
| hsa-miR-200b-3p | 2899.3 | 1172.11 | -1.3066 | down | 6.23E-05 | 0.002528 |
| hsa-miR-92b-5p | 11.67333333 | 4.766666667 | -1.29216 | down | 0.001251 | 0.025961 |
| hsa-miR-301b-3p | 18.27 | 7.5 | -1.28451 | down | 0.006431 | 0.083157 |
| hsa-miR-182-5p | 524.1166667 | 216.19 | -1.27759 | down | 0.001163 | 0.024738 |
| hsa-miR-6507-5p | 5.7 | 2.356666667 | -1.27421 | down | 0.011751 | 0.128271 |
| hsa-miR-10a-3p | 84.01333333 | 34.75333333 | -1.27347 | down | 0.000271 | 0.007734 |
| hsa-miR-130b-3p | 94.68 | 39.21333333 | -1.27172 | down | 0.00164 | 0.031195 |
| hsa-miR-486-5p | 488.06 | 203.3833333 | -1.26286 | down | 0.038294 | 0.302216 |
| hsa-miR-224-3p | 89.34666667 | 37.28 | -1.26101 | down | 0.000243 | 0.0073 |
| hsa-miR-1270 | 4.593333333 | 1.916666667 | -1.26094 | down | 0.018817 | 0.181507 |
| hsa-miR-188-5p | 69.80666667 | 29.14333333 | -1.2602 | down | 0.001401 | 0.028037 |
| hsa-miR-200a-3p | 4187.353333 | 1763.513333 | -1.24759 | down | 5.11E-05 | 0.002271 |
| hsa-miR-146b-3p | 30.36666667 | 12.8 | -1.24634 | down | 0.001917 | 0.035023 |
| hsa-miR-597-5p | 6.013333333 | 2.553333333 | -1.23578 | down | 0.012088 | 0.129096 |
| hsa-miR-21-3p | 548.0533333 | 233.11 | -1.23331 | down | 0.00302 | 0.047444 |
| hsa-miR-766-3p | 11.32 | 4.82 | -1.23177 | down | 0.002534 | 0.042179 |
| hsa-miR-181b-2-3p | 6.346666667 | 2.716666667 | -1.22416 | down | 0.006669 | 0.084092 |
| hsa-miR-19a-5p | 1.59 | 0.69 | -1.20436 | down | 0.042766 | 0.308277 |
| hsa-miR-18a-5p | 190.82 | 83.20333333 | -1.1975 | down | 0.00134 | 0.027149 |
| hsa-miR-452-3p | 64.94 | 28.42333333 | -1.19203 | down | 0.000574 | 0.014405 |
| hsa-miR-106b-5p | 2891.57 | 1265.82 | -1.19178 | down | 0.000214 | 0.006692 |
| hsa-miR-93-5p | 2033.6 | 891.6866667 | -1.18943 | down | 0.000252 | 0.007397 |
| hsa-miR-323a-3p | 43.58 | 19.37333333 | -1.16959 | down | 0.004914 | 0.065641 |
| hsa-miR-96-5p | 432.56 | 193.4 | -1.16131 | down | 0.000722 | 0.016637 |
| hsa-miR-182-3p | 4.666666667 | 2.086666667 | -1.16119 | down | 0.017348 | 0.173584 |
| hsa-miR-203a-3p | 170.3733333 | 76.48 | -1.15555 | down | 0.001097 | 0.02364 |
| hsa-miR-146b-5p | 5472.2 | 2458.666667 | -1.15424 | down | 0.002602 | 0.042878 |
| hsa-miR-21-5p | 154716.4267 | 69869.23667 | -1.1469 | down | 0.002716 | 0.044325 |
| hsa-miR-92a-3p | 8268.91 | 3740.263333 | -1.14456 | down | 0.000166 | 0.005684 |
| hsa-miR-1197 | 8.18 | 3.703333333 | -1.14328 | down | 0.026433 | 0.236352 |
| hsa-miR-767-5p | 7.796666667 | 3.536666667 | -1.14047 | down | 0.020895 | 0.192992 |
| hsa-miR-17-5p | 2417.473333 | 1102.613333 | -1.13257 | down | 0.00059 | 0.014539 |
| hsa-miR-452-5p | 180.32 | 83.15 | -1.11677 | down | 0.000678 | 0.016046 |
| hsa-miR-210-5p | 2.233333333 | 1.046666667 | -1.0934 | down | 0.04083 | 0.305049 |
| hsa-miR-1307-3p | 117.66 | 55.53666667 | -1.08311 | down | 0.001788 | 0.033388 |
| hsa-miR-181c-3p | 37.90666667 | 18.22666667 | -1.0564 | down | 0.011703 | 0.128271 |
| hsa-miR-20a-5p | 2907.733333 | 1400.2 | -1.05426 | down | 0.000769 | 0.016797 |
| hsa-miR-942-5p | 4.35 | 2.1 | -1.05063 | down | 0.036492 | 0.290726 |
| hsa-miR-32-5p | 625.2333333 | 306.22 | -1.02983 | down | 0.004114 | 0.059314 |
| hsa-miR-3173-5p | 2.04 | 1.003333333 | -1.02377 | down | 0.045708 | 0.316192 |
| hsa-miR-425-5p | 369.4933333 | 181.86 | -1.02272 | down | 0.004356 | 0.06051 |
| hsa-miR-200c-3p | 2863.576667 | 1415.84 | -1.01616 | down | 0.0038 | 0.056523 |
| hsa-miR-550a-5p | 3.76 | 1.866666667 | -1.01027 | down | 0.049216 | 0.336241 |
