## Supplementary File 2 for "Targeting circPTPN12/miR-21-5p/ΔNp63α pathway as a therapeutic strategy for human endometrial fibrosis"

Supplementary File 2. Forty upregulated and forty downregulated miRNAs with higher expression abundance.

| GeneName | Con_TPM | IUA_TPM | log2(IUA/Con) | up-or-down | p_value | q_value |
| --- | --- | --- | --- | --- | --- | --- |
| hsa-miR-21-5p | 154716.4 | 69869.24 | -1.1469 | down | 0.002716 | 0.044325 |
| hsa-miR-143-3p | 25185.46 | 116029.7 | 2.203831 | up | 1.03E-12 | 2.47E-10 |
| hsa-miR-10b-5p | 16433.64 | 36829.03 | 1.164191 | up | 0.000109 | 0.004279 |
| hsa-miR-451a | 15701.32 | 6017.757 | -1.38359 | down | 0.027449 | 0.241385 |
| hsa-miR-424-5p | 10544.78 | 24695.72 | 1.227731 | up | 0.000569 | 0.014405 |
| hsa-miR-92a-3p | 8268.91 | 3740.263 | -1.14456 | down | 0.000166 | 0.005684 |
| hsa-miR-146b-5p | 5472.2 | 2458.667 | -1.15424 | down | 0.002602 | 0.042878 |
| hsa-miR-200a-3p | 4187.353 | 1763.513 | -1.24759 | down | 5.11E-05 | 0.002271 |
| hsa-miR-20a-5p | 2907.733 | 1400.2 | -1.05426 | down | 0.000769 | 0.016797 |
| hsa-miR-200b-3p | 2899.3 | 1172.11 | -1.3066 | down | 6.23E-05 | 0.002528 |
| hsa-miR-106b-5p | 2891.57 | 1265.82 | -1.19178 | down | 0.000214 | 0.006692 |
| hsa-miR-200c-3p | 2863.577 | 1415.84 | -1.01616 | down | 0.0038 | 0.056523 |
| hsa-miR-17-5p | 2417.473 | 1102.613 | -1.13257 | down | 0.00059 | 0.014539 |
| hsa-miR-144-3p | 2204.85 | 723.12 | -1.60837 | down | 0.017532 | 0.173732 |
| hsa-miR-145-5p | 2105.527 | 9658.62 | 2.197636 | up | 1.23E-10 | 2.58E-08 |
| hsa-miR-93-5p | 2033.6 | 891.6867 | -1.18943 | down | 0.000252 | 0.007397 |
| hsa-miR-449a | 1599.377 | 242.3267 | -2.72248 | down | 1.46E-09 | 2.45E-07 |
| hsa-miR-92b-3p | 1062.617 | 353.7533 | -1.58681 | down | 1.36E-05 | 0.000844 |
| hsa-miR-429 | 809.0567 | 321.21 | -1.33272 | down | 5.74E-05 | 0.002414 |
| hsa-miR-29c-3p | 758.97 | 1627.207 | 1.100283 | up | 0.004332 | 0.06051 |
| hsa-miR-145-3p | 702.35 | 2125.387 | 1.597463 | up | 1.92E-06 | 0.000161 |
| hsa-miR-32-5p | 625.2333 | 306.22 | -1.02983 | down | 0.004114 | 0.059314 |
| hsa-miR-21-3p | 548.0533 | 233.11 | -1.23331 | down | 0.00302 | 0.047444 |
| hsa-miR-182-5p | 524.1167 | 216.19 | -1.27759 | down | 0.001163 | 0.024738 |
| hsa-miR-143-5p | 511.2267 | 1639.493 | 1.681215 | up | 8.83E-07 | 7.81E-05 |
| hsa-miR-486-5p | 488.06 | 203.3833 | -1.26286 | down | 0.038294 | 0.302216 |
| hsa-miR-96-5p | 432.56 | 193.4 | -1.16131 | down | 0.000722 | 0.016637 |
| hsa-miR-449c-5p | 414.4967 | 64.27333 | -2.68907 | down | 3.76E-08 | 4.86E-06 |
| hsa-miR-31-5p | 405.3567 | 128.7433 | -1.65469 | down | 2.09E-05 | 0.001132 |
| hsa-miR-425-5p | 369.4933 | 181.86 | -1.02272 | down | 0.004356 | 0.06051 |
| hsa-miR-28-5p | 321.2733 | 717.3133 | 1.158802 | up | 0.000736 | 0.016712 |
| hsa-miR-375 | 272.08 | 566.67 | 1.058478 | up | 0.028423 | 0.246282 |
| hsa-miR-28-3p | 225.3833 | 509.0467 | 1.175417 | up | 0.000301 | 0.008287 |
| hsa-miR-135b-5p | 215.7667 | 43.15333 | -2.32193 | down | 5.98E-09 | 8.38E-07 |
| hsa-miR-18a-5p | 190.82 | 83.20333 | -1.1975 | down | 0.00134 | 0.027149 |
| hsa-miR-452-5p | 180.32 | 83.15 | -1.11677 | down | 0.000678 | 0.016046 |
| hsa-miR-203a-3p | 170.3733 | 76.48 | -1.15555 | down | 0.001097 | 0.02364 |
| hsa-miR-204-5p | 168.9967 | 388.7533 | 1.20186 | up | 0.009568 | 0.109956 |
| hsa-miR-183-5p | 152.2433 | 41.13333 | -1.888 | down | 3.72E-06 | 0.000284 |
| hsa-miR-1307-3p | 117.66 | 55.53667 | -1.08311 | down | 0.001788 | 0.033388 |
| hsa-miR-130b-5p | 114.7267 | 31.20333 | -1.87843 | down | 5.13E-05 | 0.002271 |
| hsa-miR-130b-3p | 94.68 | 39.21333 | -1.27172 | down | 0.00164 | 0.031195 |
| hsa-miR-224-3p | 89.34667 | 37.28 | -1.26101 | down | 0.000243 | 0.0073 |
| hsa-miR-10a-3p | 84.01333 | 34.75333 | -1.27347 | down | 0.000271 | 0.007734 |
| hsa-miR-320b | 80.28333 | 185.6567 | 1.209465 | up | 0.016744 | 0.169558 |
| hsa-miR-205-5p | 76.63667 | 13.00667 | -2.55878 | down | 3.17E-05 | 0.001614 |
| hsa-miR-188-5p | 69.80667 | 29.14333 | -1.2602 | down | 0.001401 | 0.028037 |
| hsa-miR-15b-3p | 69.58667 | 22.18667 | -1.64912 | down | 0.000232 | 0.007087 |
| hsa-miR-452-3p | 64.94 | 28.42333 | -1.19203 | down | 0.000574 | 0.014405 |
| hsa-miR-4286 | 60.04 | 22.97667 | -1.38575 | down | 0.000189 | 0.006233 |
| hsa-miR-31-3p | 51.75 | 10.67333 | -2.27755 | down | 6.65E-06 | 0.000486 |
| hsa-miR-124-3p | 50.46 | 2.773333 | -4.18545 | down | 1.11E-05 | 0.000718 |
| hsa-miR-1247-5p | 32.18 | 85.14667 | 1.403786 | up | 0.000654 | 0.015712 |
| hsa-miR-1-3p | 25.51333 | 2035.3 | 6.317846 | up | 1.76E-38 | 2.96E-35 |
| hsa-miR-7641 | 21.24667 | 64.08667 | 1.592788 | up | 0.03977 | 0.304609 |
| hsa-miR-320c | 20.43667 | 83.01333 | 2.022183 | up | 0.004203 | 0.059379 |
| hsa-miR-133a-3p | 14.65667 | 1150.85 | 6.294999 | up | 6.73E-38 | 5.66E-35 |
| hsa-miR-873-5p | 11.28333 | 32.51333 | 1.526838 | up | 3.27E-05 | 0.001617 |
| hsa-miR-320d | 10.66 | 43.35333 | 2.023935 | up | 0.005635 | 0.074592 |
| hsa-miR-137 | 9.03 | 30.15667 | 1.739679 | up | 0.012557 | 0.131927 |
| hsa-miR-885-5p | 8.08 | 45.11667 | 2.481233 | up | 6.78E-08 | 7.60E-06 |
| hsa-miR-196a-5p | 7.746667 | 46.44 | 2.58372 | up | 3.92E-05 | 0.001829 |
| hsa-miR-1248 | 6.536667 | 38.54 | 2.55973 | up | 0.002011 | 0.036345 |
| hsa-miR-346 | 1.886667 | 4.203333 | 1.155694 | up | 0.028723 | 0.247388 |
| hsa-miR-1179 | 1.426667 | 4.62 | 1.695245 | up | 0.004776 | 0.064741 |
| hsa-miR-585-3p | 1.24 | 3.456667 | 1.479041 | up | 0.043589 | 0.308277 |
| hsa-miR-202-5p | 1.033333 | 3.046667 | 1.559926 | up | 0.01196 | 0.128879 |
| hsa-miR-873-3p | 0.973333 | 3.15 | 1.694346 | up | 0.001894 | 0.034994 |
| hsa-miR-133b | 0.893333 | 92.96 | 6.701268 | up | 5.48E-31 | 3.07E-28 |
| hsa-miR-6087 | 0.886667 | 2.916667 | 1.717857 | up | 0.049997 | 0.338891 |
| hsa-miR-544a | 0.69 | 1.543333 | 1.161381 | up | 0.043647 | 0.308277 |
| hsa-miR-3154 | 0.596667 | 1.406667 | 1.237283 | up | 0.040035 | 0.304609 |
| hsa-miR-490-3p | 0.563333 | 5.17 | 3.198104 | up | 4.47E-07 | 4.18E-05 |
| hsa-miR-4449 | 0.49 | 3.793333 | 2.952612 | up | 0.004147 | 0.059314 |
| hsa-miR-5683 | 0.47 | 1.443333 | 1.618672 | up | 0.013343 | 0.138458 |
| hsa-miR-619-5p | 0.436667 | 1.853333 | 2.085518 | up | 0.008945 | 0.105151 |
| hsa-miR-605-3p | 0.406667 | 2.053333 | 2.336049 | up | 0.00014 | 0.005008 |
| hsa-miR-551b-3p | 0.406667 | 1.586667 | 1.96408 | up | 0.002933 | 0.046512 |
| hsa-miR-3651 | 0.376667 | 1.48 | 1.974237 | up | 0.044453 | 0.311356 |
| hsa-miR-133a-5p | 0.266667 | 30.26333 | 6.82639 | up | 4.75E-26 | 2.00E-23 |
