## Supplementary File 3 for "Targeting circPTPN12/miR-21-5p/ΔNp63α pathway as a therapeutic strategy for human endometrial fibrosis"

Supplementary File 3. circRNAs with binding sites for miR-21-5p.

| Row data ID | Circbase ID | Gene Location | Con_TPM | IUA_  TPM | log2(IUA/Con) | up-or-down | p_value |
| --- | --- | --- | --- | --- | --- | --- | --- |
| hsa_circRNA031909 | hsa_circ_0003764 | PTPN12 | 0.001 | 374.09 | 9.228615 | up | 0.002264 |
| hsa_circRNA036947 | hsa_circ_0006764 | MTMR1 | 0.001 | 507.8933 | 9.669584 | up | 0.006272 |
| hsa_circRNA013374 | hsa_circ_0039550 | RSPRY1 | 323.6967 | 0.001 | -9.15644 | down | 0.009906 |
| hsa_circRNA004522 | hsa_circ_0008629 | STAMBPL1 | 316.1167 | 0.001 | -9.11866 | down | 0.010759 |
| hsa_circRNA029894 | hsa_circ_0076961 | FAM135A | 0.001 | 314.86 | 8.973013 | up | 0.01141 |
| hsa_circRNA026069 | hsa_circ_0003936 | EXOC1 | 0.001 | 293.9267 | 8.874602 | up | 0.012485 |
| hsa_circRNA008473 | hsa_circ_0006512 | RIC8B | 0.001 | 269.8433 | 8.771145 | up | 0.01393 |
| hsa_circRNA019140 | hsa_circ_0054958 | VPS54 | 0.001 | 276.3267 | 8.778152 | up | 0.01431 |
| hsa_circRNA032345 | hsa_circ_0006252 | CADPS2 | 0.001 | 199.0967 | 8.312265 | up | 0.021559 |
| hsa_circRNA011775 | hsa_circ_0035446 | ALDH1A2 | 171.3367 | 0.001 | -8.23751 | down | 0.024156 |
| hsa_circRNA027149 | hsa_circ_0005274 | DCTD | 152.7267 | 0.001 | -8.06069 | down | 0.028422 |
| hsa_circRNA000676 | hsa_circ_0011162 | TAF12 | 0.001 | 154.4133 | 7.962349 | up | 0.029515 |
| hsa_circRNA027134 | hsa_circ_0004160 | VEGFC | 0.001 | 132.8733 | 7.740243 | up | 0.036107 |
| hsa_circRNA016367 | hsa_circ_0047870 | KIAA1468 | 0.001 | 125.18 | 7.646458 | up | 0.039244 |
| hsa_circRNA033398 | hsa_circ_0084501 | RB1CC1 | 0.001 | 119.7633 | 7.595837 | up | 0.041427 |
| hsa_circRNA019332 | hsa_circ_0055241 | ALMS1 | 0.001 | 107.7433 | 7.434661 | up | 0.047623 |
| hsa_circRNA009480 | hsa_circ_0007259 | GTF2F2 | 0.001 | 91.70667 | 7.202867 | up | 0.058871 |
| hsa_circRNA026767 | hsa_circ_0008825 | ELF2 | 79.46667 | 0.001 | -7.13835 | down | 0.067958 |
| hsa_circRNA036622 | hsa_circ_0091103 | ATRX | 78.00333 | 0.001 | -7.09685 | down | 0.069582 |
| hsa_circRNA021302 | hsa_circ_0001137 | DNMT3B | 73.89667 | 0.001 | -7.01881 | down | 0.072724 |
| hsa_circRNA020749 | hsa_circ_0058480 | CUL3 | 0.001 | 76.23333 | 6.937848 | up | 0.07601 |
| hsa_circRNA018591 | hsa_circ_0006294 | CRIM1 | 68.85667 | 0.001 | -6.90363 | down | 0.077573 |
| hsa_circRNA019251 | hsa_circ_0055102 | PCBP1-AS1 | 0.001 | 63.9 | 6.683111 | up | 0.087471 |
| hsa_circRNA013201 | hsa_circ_0039202 | VPS35 | 56.93333 | 0.001 | -6.65729 | down | 0.088834 |
| hsa_circRNA008345 | hsa_circ_0004901 | APAF1 | 54.42 | 0.001 | -6.56435 | down | 0.093413 |
| hsa_circRNA010447 | hsa_circ_0031970 | FERMT2 | 54.42 | 0.001 | -6.56435 | down | 0.093413 |
| hsa_circRNA030674 | hsa_circ_0078210 | C6orf72 | 53.37 | 0.001 | -6.549 | down | 0.094187 |
| hsa_circRNA022515 | hsa_circ_0002063 | FBXO7 | 52.2 | 0.001 | -6.50427 | down | 0.096472 |
| hsa_circRNA016337 | hsa_circ_0047815 | ZNF532 | 0.001 | 51.82667 | 6.396472 | up | 0.102001 |
| hsa_circRNA030650 | hsa_circ_0078165 | STXBP5 | 45.53333 | 0.001 | -6.30739 | down | 0.107069 |
| hsa_circRNA026015 | hsa_circ_0001410 | DCUN1D4 | 0.001 | 47.11667 | 6.259113 | up | 0.109618 |
| hsa_circRNA034816 | hsa_circ_0003654 | GCNT1 | 0.001 | 46.85667 | 6.212558 | up | 0.1123 |
| hsa_circRNA026481 | hsa_circ_0001431 | MANBA | 0.001 | 41.48 | 6.059121 | up | 0.121508 |
| hsa_circRNA018881 | hsa_circ_0006911 | PPP1R21 | 0.001 | 39.88 | 5.979876 | up | 0.126491 |
| hsa_circRNA034313 | hsa_circ_0005890 | GLIS3 | 0.001 | 36.90667 | 5.907245 | up | 0.131198 |
| hsa_circRNA022512 | hsa_circ_0062983 | FBXO7 | 0.001 | 37.88333 | 5.905869 | up | 0.131288 |
| hsa_circRNA011398 | hsa_circ_0000593 | MGA | 0.001 | 34.89333 | 5.78721 | up | 0.139274 |
| hsa_circRNA004493 | hsa_circ_0019014 | WAPAL | 0.001 | 31.39 | 5.656513 | up | 0.1485 |
| hsa_circRNA013203 | hsa_circ_0002500 | VPS35 | 0.001 | 29.14667 | 5.549437 | up | 0.156404 |
| hsa_circRNA028614 | hsa_circ_0003205 | CTNNA1 | 26.65333 | 0.001 | -5.53562 | down | 0.157806 |
| hsa_circRNA012050 | hsa_circ_0035999 | ZWILCH | 24.63333 | 0.001 | -5.43218 | down | 0.165817 |
| hsa_circRNA020806 | hsa_circ_0008382 | SP140L | 23.72333 | 0.001 | -5.39452 | down | 0.168809 |
| hsa_circRNA033953 | hsa_circ_0085363 | TRPS1 | 608.6133 | 24.66333 | -4.51921 | down | 0.170232 |
| hsa_circRNA016309 | hsa_circ_0047744 | WDR7 | 0.001 | 23.55667 | 5.260669 | up | 0.17932 |
| hsa_circRNA026359 | hsa_circ_0070389 | KLHL8 | 865.0933 | 115.5233 | -2.77625 | down | 0.28423 |
| hsa_circRNA020615 | hsa_circ_0058010 | C2orf67 | 919.64 | 82.96 | -3.36615 | down | 0.317639 |
| hsa_circRNA007872 | hsa_circ_0026235 | LARP4 | 26.65333 | 240.5933 | 3.294912 | up | 0.322231 |
| hsa_circRNA020621 | hsa_circ_0008459 | C2orf67 | 1273.427 | 3756.707 | 1.667322 | up | 0.423448 |
| hsa_circRNA019334 | hsa_circ_0055243 | ALMS1 | 730.7233 | 133.4067 | -2.34489 | down | 0.427799 |
| hsa_circRNA022621 | hsa_circ_0001231 | DMC1 | 267.84 | 39.88 | -2.65499 | down | 0.429803 |
| hsa_circRNA011646 | hsa_circ_0035265 | SPPL2A | 251.5333 | 47.11667 | -2.29605 | down | 0.495965 |
| hsa_circRNA008807 | hsa_circ_0029069 | CLIP1 | 133.33 | 393.6633 | 1.659534 | up | 0.516243 |
| hsa_circRNA026016 | hsa_circ_0007646 | DCUN1D4 | 367.2433 | 109.6667 | -1.6269 | down | 0.528823 |
| hsa_circRNA018348 | hsa_circ_0000983 | NCOA1 | 473.1867 | 104.68 | -2.09185 | down | 0.536101 |
| hsa_circRNA007859 | hsa_circ_0026228 | LARP4 | 105.5067 | 348.85 | 1.848387 | up | 0.583528 |
| hsa_circRNA012294 | hsa_circ_0005558 | MESDC2 | 233.5567 | 539.2967 | 1.298417 | up | 0.620916 |
| hsa_circRNA036984 | hsa_circ_0006355 | TMLHE | 239.48 | 612.1567 | 1.470022 | up | 0.624953 |
| hsa_circRNA004818 | hsa_circ_0005620 | SH3PXD2A | 1673.037 | 758.5 | -1.03434 | down | 0.627499 |
| hsa_circRNA030466 | hsa_circ_0001640 | EPB41L2 | 326.2567 | 723.0933 | 1.25943 | up | 0.630685 |
| hsa_circRNA002924 | hsa_circ_0004417 | LYPLAL1 | 746.7033 | 1385.173 | 0.996245 | up | 0.633877 |
| hsa_circRNA030465 | hsa_circ_0077837 | EPB41L2 | 424.3167 | 800.59 | 1.031881 | up | 0.635985 |
| hsa_circRNA015399 | hsa_circ_0006479 | TEX2 | 132.1633 | 292.7267 | 1.268934 | up | 0.6692 |
| hsa_circRNA020399 | hsa_circ_0004191 | PGAP1 | 430.0833 | 217.7167 | -0.87863 | down | 0.685803 |
| hsa_circRNA005582 | hsa_circ_0021572 | QSER1 | 117.4233 | 273.04 | 1.296022 | up | 0.704477 |
| hsa_circRNA006492 | hsa_circ_0000344 | RSF1 | 789.2433 | 381.63 | -0.94202 | down | 0.720409 |
| hsa_circRNA003488 | hsa_circ_0000209 | FAM208B | 469.3567 | 713.8867 | 0.726558 | up | 0.735546 |
| hsa_circRNA035282 | hsa_circ_0087961 | LPAR1 | 423.3833 | 218.8267 | -0.83102 | down | 0.748671 |
| hsa_circRNA000339 | hsa_circ_0010029 | PRDM2 | 209.7567 | 374.8333 | 0.956874 | up | 0.749285 |
| hsa_circRNA009091 | hsa_circ_0029625 | ZMYM2 | 394.59 | 194.8933 | -0.89629 | down | 0.764324 |
| hsa_circRNA026361 | hsa_circ_0006866 | KLHL8 | 849.8267 | 517.81 | -0.60266 | down | 0.777765 |
| hsa_circRNA000729 | hsa_circ_0000043 | PUM1 | 197.8067 | 119.35 | -0.62005 | down | 0.782255 |
| hsa_circRNA030098 | hsa_circ_0003814 | MDN1 | 265.6733 | 423.74 | 0.767706 | up | 0.798591 |
| hsa_circRNA009094 | hsa_circ_0005966 | ZMYM2 | 413.9267 | 254.9167 | -0.59854 | down | 0.816484 |
| hsa_circRNA033186 | hsa_circ_0083902 | FUT10 | 95.79333 | 143.5767 | 0.685874 | up | 0.820727 |
| hsa_circRNA031904 | hsa_circ_0080835 | PTPN12 | 173.8667 | 231.1467 | 0.52234 | up | 0.839103 |
| hsa_circRNA034125 | hsa_circ_0007209 | ZFAT | 62.66667 | 35.89 | -0.71365 | down | 0.841747 |
| hsa_circRNA010783 | hsa_circ_0032649 | MLH3 | 3253.703 | 2262.253 | -0.41098 | down | 0.843273 |
| hsa_circRNA003844 | hsa_circ_0000230 | ZEB1 | 587.1467 | 404.97 | -0.42131 | down | 0.844252 |
| hsa_circRNA009703 | hsa_circ_0000497 | SLAIN1 | 691.7467 | 486.4833 | -0.41563 | down | 0.84948 |
| hsa_circRNA026017 | hsa_circ_0069718 | DCUN1D4 | 195.0367 | 123.84 | -0.5601 | down | 0.852322 |
| hsa_circRNA023671 | hsa_circ_0001306 | RAD54L2 | 702.21 | 472.7933 | -0.46309 | down | 0.858865 |
| hsa_circRNA011529 | hsa_circ_0003784 | SPG11 | 26.65333 | 38.88333 | 0.645286 | up | 0.863192 |
| hsa_circRNA016335 | hsa_circ_0047814 | ZNF532 | 1678.417 | 1225.447 | -0.33974 | down | 0.872761 |
| hsa_circRNA035278 | hsa_circ_0004928 | LPAR1 | 258.4167 | 171.4233 | -0.47729 | down | 0.874391 |
| hsa_circRNA020097 | hsa_circ_0002603 | UBR3 | 138.8267 | 180.91 | 0.51737 | up | 0.87961 |
| hsa_circRNA022511 | hsa_circ_0001222 | FBXO7 | 311.6 | 236.3 | -0.30328 | down | 0.890469 |
| hsa_circRNA002214 | hsa_circ_0014591 | ASH1L | 187 | 228.2467 | 0.412911 | up | 0.891174 |
| hsa_circRNA004124 | hsa_circ_0005144 | JMJD1C | 173.1567 | 215.3167 | 0.406575 | up | 0.892439 |
| hsa_circRNA011772 | hsa_circ_0035443 | ALDH1A2 | 720.2467 | 789.0467 | 0.254944 | up | 0.906085 |
| hsa_circRNA007588 | hsa_circ_0025763 | TMTC1 | 73.53667 | 50.25667 | -0.43679 | down | 0.9069 |
| hsa_circRNA014480 | hsa_circ_0008603 | ALDH3A2 | 355.6533 | 271.83 | -0.29762 | down | 0.921436 |
| hsa_circRNA033949 | hsa_circ_0085362 | TRPS1 | 1401.9 | 1073.687 | -0.2576 | down | 0.921727 |
| hsa_circRNA028615 | hsa_circ_0007440 | CTNNA1 | 477.8233 | 379.8867 | -0.21481 | down | 0.933573 |
| hsa_circRNA021087 | hsa_circ_0001127 | PTPRA | 149.4333 | 158.66 | 0.180472 | up | 0.952213 |
| hsa_circRNA018649 | hsa_circ_0000992 | PRKD3 | 2313.74 | 2238.53 | 0.054258 | up | 0.979738 |
| hsa_circRNA009577 | hsa_circ_0005263 | FNDC3A | 321.2367 | 287.24 | -0.05664 | down | 0.985099 |
| hsa_circRNA020618 | hsa_circ_0002617 | C2orf67 | 845.3833 | 797.5633 | 0.023155 | up | 0.991348 |
| hsa_circRNA020620 | hsa_circ_0002822 | C2orf67 | 311.2267 | 288.9067 | -0.00768 | down | 0.997645 |
| hsa_circRNA036804 | hsa_circ_0006727 | SLC25A43 | 0.001 | 493.14 | 23.63306 | up | 1 |
| hsa_circRNA036803 | hsa_circ_0005213 | SLC25A43 | 0.001 | 279.1433 | 22.84664 | up | 1 |
| hsa_circRNA009807 | hsa_circ_0030716 | DOCK9 | 0.001 | 228.6967 | 22.57472 | up | 1 |
| hsa_circRNA011047 | hsa_circ_0033184 | WARS | 0.001 | 213.0033 | 22.47374 | up | 1 |
| hsa_circRNA019118 | hsa_circ_0007387 | WDPCP | 0.001 | 202.3833 | 22.38768 | up | 1 |
| hsa_circRNA019648 | hsa_circ_0003523 | SH3RF3 | 0.001 | 188.34 | 22.31088 | up | 1 |
| hsa_circRNA034454 | hsa_circ_0003841 | BNC2 | 0.001 | 186.1067 | 22.31032 | up | 1 |
| hsa_circRNA025206 | hsa_circ_0005494 | ATP13A3 | 0.001 | 176.6833 | 22.23866 | up | 1 |
| hsa_circRNA019803 | hsa_circ_0056426 | UGGT1 | 0.001 | 172.7567 | 22.20254 | up | 1 |
| hsa_circRNA019133 | hsa_circ_0054951 | VPS54 | 0.001 | 170.4 | 22.17375 | up | 1 |
| hsa_circRNA005709 | hsa_circ_0021902 | CKAP5 | 0.001 | 166.4933 | 22.12005 | up | 1 |
| hsa_circRNA014670 | hsa_circ_0042881 | NF1 | 0.001 | 138.99 | 21.9136 | up | 1 |
| hsa_circRNA016307 | hsa_circ_0006090 | WDR7 | 0.001 | 155.5267 | 21.8753 | up | 1 |
| hsa_circRNA019252 | hsa_circ_0055103 | PCBP1-AS1 | 0.001 | 124.07 | 21.76007 | up | 1 |
| hsa_circRNA015241 | hsa_circ_0004090 | TRIM37 | 0.001 | 123.2833 | 21.75149 | up | 1 |
| hsa_circRNA007862 | hsa_circ_0026230 | LARP4 | 0.001 | 114.35 | 21.63513 | up | 1 |
| hsa_circRNA030823 | hsa_circ_0001658 | ARID1B | 0.001 | 109.6667 | 21.55754 | up | 1 |
| hsa_circRNA035690 | hsa_circ_0089277 | SETX | 0.001 | 107.6233 | 21.55545 | up | 1 |
| hsa_circRNA023729 | hsa_circ_0001974 | TKT | 0.001 | 106.01 | 21.54678 | up | 1 |
| hsa_circRNA004014 | hsa_circ_0002144 | MAPK8 | 0.001 | 99.77333 | 21.45358 | up | 1 |
| hsa_circRNA015391 | hsa_circ_0045226 | DDX42 | 0.001 | 94.23 | 21.38969 | up | 1 |
| hsa_circRNA008968 | hsa_circ_0029408 | GLT1D1 | 0.001 | 91.09 | 21.15346 | up | 1 |
| hsa_circRNA025024 | hsa_circ_0068045 | TBL1XR1 | 0.001 | 91.09 | 21.15346 | up | 1 |
| hsa_circRNA031916 | hsa_circ_0080865 | RSBN1L | 0.001 | 85.59333 | 7.11969 | up | 1 |
| hsa_circRNA005545 | hsa_circ_0007276 | METTL15 | 0.001 | 87.73333 | 7.117451 | up | 1 |
| hsa_circRNA010094 | hsa_circ_0006038 | STXBP6 | 0.001 | 81.66667 | 7.051988 | up | 1 |
| hsa_circRNA016792 | hsa_circ_0002866 | ZNF562 | 0.001 | 81.66667 | 7.051988 | up | 1 |
| hsa_circRNA016338 | hsa_circ_0047816 | ZNF532 | 0.001 | 81.83667 | 7.040268 | up | 1 |
| hsa_circRNA022514 | hsa_circ_0062984 | FBXO7 | 0.001 | 75.38333 | 6.936588 | up | 1 |
| hsa_circRNA003618 | hsa_circ_0004201 | DCLRE1C | 0.001 | 73.81333 | 6.906236 | up | 1 |
| hsa_circRNA018417 | hsa_circ_0000985 | ZNF512 | 0.001 | 74.77333 | 6.886829 | up | 1 |
| hsa_circRNA019534 | hsa_circ_0004147 | MRPL30 | 0.001 | 69.88667 | 6.827438 | up | 1 |
| hsa_circRNA027551 | hsa_circ_0072437 | PARP8 | 0.001 | 69.10333 | 6.811148 | up | 1 |
| hsa_circRNA015520 | hsa_circ_0004716 | WIPI1 | 0.001 | 62.82 | 6.673754 | up | 1 |
| hsa_circRNA022088 | hsa_circ_0061766 | WRB | 0.001 | 56.53667 | 6.521883 | up | 1 |
| hsa_circRNA029033 | hsa_circ_0002972 | FAM193B | 98.84333 | 344.1667 | 1.925732 | up | 1 |
| hsa_circRNA006409 | hsa_circ_0023555 | C2CD3 | 72.35 | 196.4033 | 1.514635 | up | 1 |
| hsa_circRNA010788 | hsa_circ_0003670 | NEK9 | 47.44333 | 71.78333 | 0.671203 | up | 1 |
| hsa_circRNA000997 | hsa_circ_0009142 | CAP1 | 99.89667 | 117.7867 | 0.363691 | up | 1 |
| hsa_circRNA035277 | hsa_circ_0002287 | LPAR1 | 111.06 | 126.68 | 0.315472 | up | 1 |
| hsa_circRNA008471 | hsa_circ_0008366 | RIC8B | 143.2667 | 159.19 | 0.277886 | up | 1 |
| hsa_circRNA021668 | hsa_circ_0060735 | CSE1L | 157.7067 | 172.6433 | 0.256468 | up | 1 |
| hsa_circRNA009677 | hsa_circ_0002917 | TBC1D4 | 81.84 | 88.73 | 0.19053 | up | 1 |
| hsa_circRNA033774 | hsa_circ_0005934 | STK3 | 90.14333 | 85.74 | 0.001669 | up | 1 |
| hsa_circRNA030792 | hsa_circ_0078357 | SCAF8 | 126.9133 | 89.68333 | -0.40367 | down | 1 |
| hsa_circRNA009372 | hsa_circ_0030045 | SLC25A15 | 149.4467 | 104.68 | -0.43965 | down | 1 |
| hsa_circRNA009160 | hsa_circ_0003408 | SACS | 97.73333 | 64.80333 | -0.4907 | down | 1 |
| hsa_circRNA020344 | hsa_circ_0005503 | ANKAR | 214.6833 | 116.2167 | -0.77372 | down | 1 |
| hsa_circRNA018349 | hsa_circ_0005968 | NCOA1 | 92.18 | 46.33 | -0.85182 | down | 1 |
| hsa_circRNA016310 | hsa_circ_0047745 | WDR7 | 209.9033 | 99.72667 | -0.93352 | down | 1 |
| hsa_circRNA018597 | hsa_circ_0008959 | CRIM1 | 171.0333 | 50.84667 | -1.64778 | down | 1 |
| hsa_circRNA025833 | hsa_circ_0069399 | ARAP2 | 802.43 | 106.7933 | -2.78352 | down | 1 |
| hsa_circRNA025832 | hsa_circ_0069397 | ARAP2 | 91.07 | 0.001 | -7.30679 | down | 1 |
| hsa_circRNA004653 | hsa_circ_0006198 | LCOR | 92.51667 | 0.001 | -7.35765 | down | 1 |
| hsa_circRNA027150 | hsa_circ_0002212 | DCTD | 111.06 | 0.001 | -19.1788 | down | 1 |
| hsa_circRNA019125 | hsa_circ_0006759 | WDPCP | 135.2133 | 0.001 | -19.7269 | down | 1 |
| hsa_circRNA006490 | hsa_circ_0023700 | RSF1 | 130.0033 | 0.001 | -20.2421 | down | 1 |
| hsa_circRNA035697 | hsa_circ_0001898 | SETX | 102.1767 | 0.001 | -20.8299 | down | 1 |
| hsa_circRNA024677 | hsa_circ_0067599 | GK5 | 102.0033 | 0.001 | -20.8528 | down | 1 |
| hsa_circRNA006491 | hsa_circ_0023701 | RSF1 | 105.5633 | 0.001 | -20.9028 | down | 1 |
| hsa_circRNA028599 | hsa_circ_0006616 | ETF1 | 109.95 | 0.001 | -20.9315 | down | 1 |
| hsa_circRNA008687 | hsa_circ_0002175 | MED13L | 110.8467 | 0.001 | -20.9562 | down | 1 |
| hsa_circRNA026306 | hsa_circ_0002283 | WDFY3 | 128.83 | 0.001 | -21.1523 | down | 1 |
| hsa_circRNA033681 | hsa_circ_0084890 | TMEM67 | 143.2667 | 0.001 | -21.2965 | down | 1 |
| hsa_circRNA030824 | hsa_circ_0004701 | ARID1B | 147.71 | 0.001 | -21.3376 | down | 1 |
| hsa_circRNA028611 | hsa_circ_0074159 | CTNNA1 | 156.0033 | 0.001 | -21.394 | down | 1 |
| hsa_circRNA002202 | hsa_circ_0008161 | ASH1L | 150.6333 | 0.001 | -21.3946 | down | 1 |
| hsa_circRNA027550 | hsa_circ_0072433 | PARP8 | 199.91 | 0.001 | -21.7636 | down | 1 |
| hsa_circRNA026360 | hsa_circ_0005384 | KLHL8 | 225.3567 | 0.001 | -21.9577 | down | 1 |
| hsa_circRNA009706 | hsa_circ_0000498 | RNF219 | 241.9633 | 0.001 | -22.0555 | down | 1 |
| hsa_circRNA008474 | hsa_circ_0028021 | RIC8B | 255.9033 | 0.001 | -22.1211 | down | 1 |
| hsa_circRNA011774 | hsa_circ_0035445 | ALDH1A2 | 279.9167 | 0.001 | -22.2595 | down | 1 |
| hsa_circRNA018344 | hsa_circ_0004938 | NCOA1 | 287.0333 | 0.001 | -22.288 | down | 1 |
| hsa_circRNA022481 | hsa_circ_0062936 | PRR14L | 303.7967 | 0.001 | -22.3587 | down | 1 |
