## Supplementary File 4 for "Targeting circPTPN12/miR-21-5p/ΔNp63α pathway as a therapeutic strategy for human endometrial fibrosis"

Supplementary File 4. Clinical information of all patients and controls.

| **Items** | **Control**  **(*n* = 18)** | **IUA**  **(*n* = 18)** | ***P* value** |
| --- | --- | --- | --- |
| Age (year) | 30.56 ±1.05 | 29.39 ± 0.95 | > 0.05 |
| Duration of infertility (year) | 1.28 ± 0.11 | 2.17 ± 0.42 | < 0.05 |
| Intrauterine adhesions under hysteroscopy (AFS score) | 0 | 9.11 ± 0.24 | < 0.05 |
| Endometrial thickness (mm)  (late-proliferative phase) | 8.67 ± 0.40 | 5.81 ± 0.35 | < 0.05 |
| Intrauterine tuberculosis or  uterine artery embolization | 0 | 0 | < 0.05 |

Note: AFS, American Fertility Society.
