## Supplementary File 5 for "Targeting circPTPN12/miR-21-5p/ΔNp63α pathway as a therapeutic strategy for human endometrial fibrosis"

Supplementary File 5. All primers used in this study.

| Gene | Species | Primer | Sequence (5’ to 3’) |
| --- | --- | --- | --- |
| miR-1304-3p | Human | Forward | ACACTCCAGCTGGGAAACtctcactgtagcct |
| miR-1304-3p | Human | Reverse | CTCAACTGGTGTCGTGGAGTCGGCAATTCAGTTGAGggggttcg |
| miR-490-5p | Human | Forward | ACACTCCAGCTGGGAAACccatggatctcc |
| miR-490-5p | Human | Reverse | CTCAACTGGTGTCGTGGAGTCGGCAATTCAGTTGAGacccacct |
| miR-7974 | Human | Forward | ACACTCCAGCTGGGAAACaggctgtgatgctctc |
| miR-7974 | Human | Reverse | CTCAACTGGTGTCGTGGAGTCGGCAATTCAGTTGAGgggctcag |
| miR-2682-5p | Human | Forward | ACACTCCAGCTGGGAAACcaggcagtgactgtt |
| miR-2682-5p | Human | Reverse | CTCAACTGGTGTCGTGGAGTCGGCAATTCAGTTGAGgacgtctg |
| miR-219a-2-3p | Human | Forward | ACACTCCAGCTGGGAAACagaattgtggctgg |
| miR-219a-2-3p | Human | Reverse | CTCAACTGGTGTCGTGGAGTCGGCAATTCAGTTGAGacagatgt |
| miR-124-5p | Human | Forward | ACACTCCAGCTGGGAAACcgtgttcacagcgg |
| miR-124-5p | Human | Reverse | CTCAACTGGTGTCGTGGAGTCGGCAATTCAGTTGAGatcaaggt |
| miR-21-5p | Human | Forward | ACACTCCAGCTGGGAAACtagcttatcagact |
| miR-21-5p | Human | Reverse | CTCAACTGGTGTCGTGGAGTCGGCAATTCAGTTGAGtcaacatc |
| URP | Human |  | TGGTGTCGTGGAGTCG |
| GAPDH | Human | Forward | GGAGCGAGATCCCTCCAAAAT |
| GAPDH | Human | Reverse | GGCTGTTGTCATACTTCTCATGG |
| 18s | Human | Forward | CTTTGGTCGCTCGCTCCTC |
| 18s | Human | Reverse | CTGACCGGGTTGGTTTTGAT |
| U6 | Human | Forward | CTCGCTTCGGCAGCACA |
| U6 | Human | Reverse | AACGCTTCACGAATTTGCGT |
| circPTPN12 (hsa_circ_0003764) | Human | Forward | TCAAAATGAATCTCGTAGGCTGT |
| circPTPN12 (hsa_circ_0003764) | Human | Reverse | GTGCAAACGTTATGGGGTCT |
| PTPN12 | Human | Forward | GATGGGAAGGAAAAAATGTGAGC |
| PTPN12 | Human | Reverse | AATGAAACTGATACAGCCTACGAGA |
| ΔNp63α | Human | Forward | GTTTCTTAGCGAGGTTGG |
| ΔNp63α | Human | Reverse | TGCTCAGGGATTTTCAG |
| E-cadherin | Human | Forward | CGAGAGCTACACGTTCACGG |
| E-cadherin | Human | Reverse | GGGTGTCGAGGGAAAAATAGG |
| N-cadherin | Human | Forward | AGCCAACCTTAACTGAGGAGT |
| N-cadherin | Human | Reverse | GGCAAGTTGATTGGAGGGATG |
| α-sma | Human | Forward | CTATGAGGGCTATGCCTTGCC |
| α-sma | Human | Reverse | GCTCAGCAGTAGTAACGAAGGA |
| FN | Human | Forward | AGGAAGCCGAGGTTTTAACTG |
| FN | Human | Reverse | AGGACGCTCATAAGTGTCACC |
| Snail | Human | Forward | TCGGAAGCCTAACTACAGCGA |
| Snail | Human | Reverse | AGATGAGCATTGGCAGCGAG |
| mus-E-cadherin | Mouse | Forward | CAGTTCCGAGGTCTACACCTT |
| mus-E-cadherin | Mouse | Reverse | TGAATCGGGAGTCTTCCGAAAA |
| mus-N-cadherin | Mouse | Forward | AGGCTTCTGGTGAAATTGCAT |
| mus-N-cadherin | Mouse | Reverse | GTCCACCTTGAAATCTGCTGG |
| mus-α-sma | Mouse | Forward | CCCAGACATCAGGGAGTAATGG |
| mus-α-sma | Mouse | Reverse | TCTATCGGATACTTCAGCGTCA |
| mus-FN | Mouse | Forward | ATGTGGACCCCTCCTGATAGT |
| mus-FN | Mouse | Reverse | GCCCAGTGATTTCAGCAAAGG |
| mus-ΔNp63α | Mouse | Forward | CATGTCACCGAGGTTGTGAAA |
| mus-ΔNp63α | Mouse | Reverse | CTGGGAGGGGCAATCTGTC |
