## Supplementary File 6 for "Targeting circPTPN12/miR-21-5p/ΔNp63α pathway as a therapeutic strategy for human endometrial fibrosis"

Supplementary File 6. Mouse model grouping.

| Treatment | Groups |
| --- | --- |
| Mechanical injury alone | Sham-operation (*n* = 3)  Mechanically injured (*n* = 3) |
| Mechanical injury + circPTPN12 overexpression | Sham-operation (*n* = 4)  Mechanically injured + AAV-control (*n* = 4)  Mechanically injured + AAV-circPTPN12 (*n* = 4) |
| miR-21-5p replenishment | Sham-operation (*n* = 3)  Mechanically injured + AAV-circPTPN12 + agomir-NC (*n* = 3)  Mechanically injured + AAV-circPTPN12 + agomir-21-5p (*n* = 3) |
